# Approach trajectory and solar position affect host plant attractiveness to the small white butterfly

**DOI:** 10.1101/2020.10.04.325639

**Authors:** Adam J. Blake, Samuel Couture, Matthew C. Go, Gerhard Gries

## Abstract

While it is well documented that insects exploit polarized sky light for navigation, their use of reflected polarized light for object detection has been less well studied. Recently, we have shown that the small white butterfly, *Pieris rapae*, distinguishes between host and non-host plants based on the degree of linear polarization (*DoLP*) of light reflected from their leaves. To determine how polarized light cues affect host plant foraging by female *P. rapae* across their entire visual range including the ultraviolet (300-650 nm), we applied photo polarimetry demonstrating large differences in the *DoLP* of leaf-reflected light among plant species generally and between host and non-host plants specifically. As polarized light cues are directionally dependent, we also tested, and modelled, the effect of approach trajectory on the polarization of plant-reflected light and the resulting attractiveness to *P. rapae*. Using photo polarimetry measurements of plants under a range of light source and observer positions, we reveal several distinct effects when polarized reflections are examined on a whole-plant basis rather than at the scale of pixels or of entire plant canopies. Most notably from our modeling, certain approach trajectories are optimal for foraging butterflies, or insects generally, to discriminate between plant species on the basis of the *DoLP* of leaf-reflected light.

## Introduction

Many insects exploit polarized skylight to aid in navigation (Labhart & Meyer, 1999) but their use of reflected polarized light for host plant detection and selection has hardly been studied (Heinloth et al., 2018). Recently, the small white butterfly, *Pieris rapae* (Linnaeus, 1758), which uses cabbage and other crucifers as host plants (Chew & Renwick, 1995), has been shown to discriminate among host and non-host plants based on the degree of linear polarization (0-100%, *DoLP*) of foliar reflections (Blake et al., 2019). Similar to many other insects (Ilić et al., 2016; Mishra, 2015; Wachmann, 1977), the rhabdom of *P. rapae* photoreceptors is untwisted with uncurved microvilli that are aligned along the rhabdom’s length (Blake et al., 2019; Qiu et al., 2002). Rhabdomeric photoreceptors have an inherent dichroism due to the tubular structure of the microvilli (Horváth & Varjú, 2004). In the ventral compound eye of other insects such as honey bees, *Apis mellifera*, desert ants, *Cataglyphis bicolor*, crickets, *Gryllus campestris*, and cockchafers, *Melolontha melolontha*, the photoreceptors along with the microvilli composing the rhabdom twist along the photoreceptor’s longitudinal axis (Wehner & Bernard, 1993). This twist serves to disrupt the alignment of microvilli along the rhabdom, preventing preferential absorption of light vibrating in a direction, or with an axis of polarization (0-180°, *AoP*), parallel to the microvillar orientation, as shown in *P. rapae* and other insects. Polarization can result in perceived shifts in color and/or intensity as compared to polarization-blind visual systems (Kelber et al., 2001; Kinoshita et al., 2011).

Shiny surfaces like water, glass or plant foliage can polarize light through specular reflection (Foster et al., 2018). These reflections are polarized in a direction so that their *AoP* is parallel to the surface. The strength of this polarization (*DoLP*) is dependent on the incident angle, with maximal polarization occurring at the Brewster’s angle (approximately 55° for foliage; Grant et al., 1993; Johnsen, 2011). The polarization of this light is consequently dependent upon the angle (ω) formed between the sun, the reflecting leaf surface, and the observer (i.e., a camera or insect; Fig. 1). This angle is itself dependent upon the solar and observer elevation and azimuth, making these aspects important predictors of foliar polarization (Hegedüs & Horváth, 2004). As it is only the specular component of the reflection that is polarized, leaf surface characteristics that increase surface roughness and diffuse reflectance, such as pubescence, epicuticular waxes or undulations, also affect the *DoLP* (Grant et al., 1993). The *DoLP* can also be altered by reducing diffuse reflectance through pigmentation absorption (Horváth & Varjú, 1997), resulting in an increased foliar *DoLP* in the red and blue relative to green.

**Figure 1.**
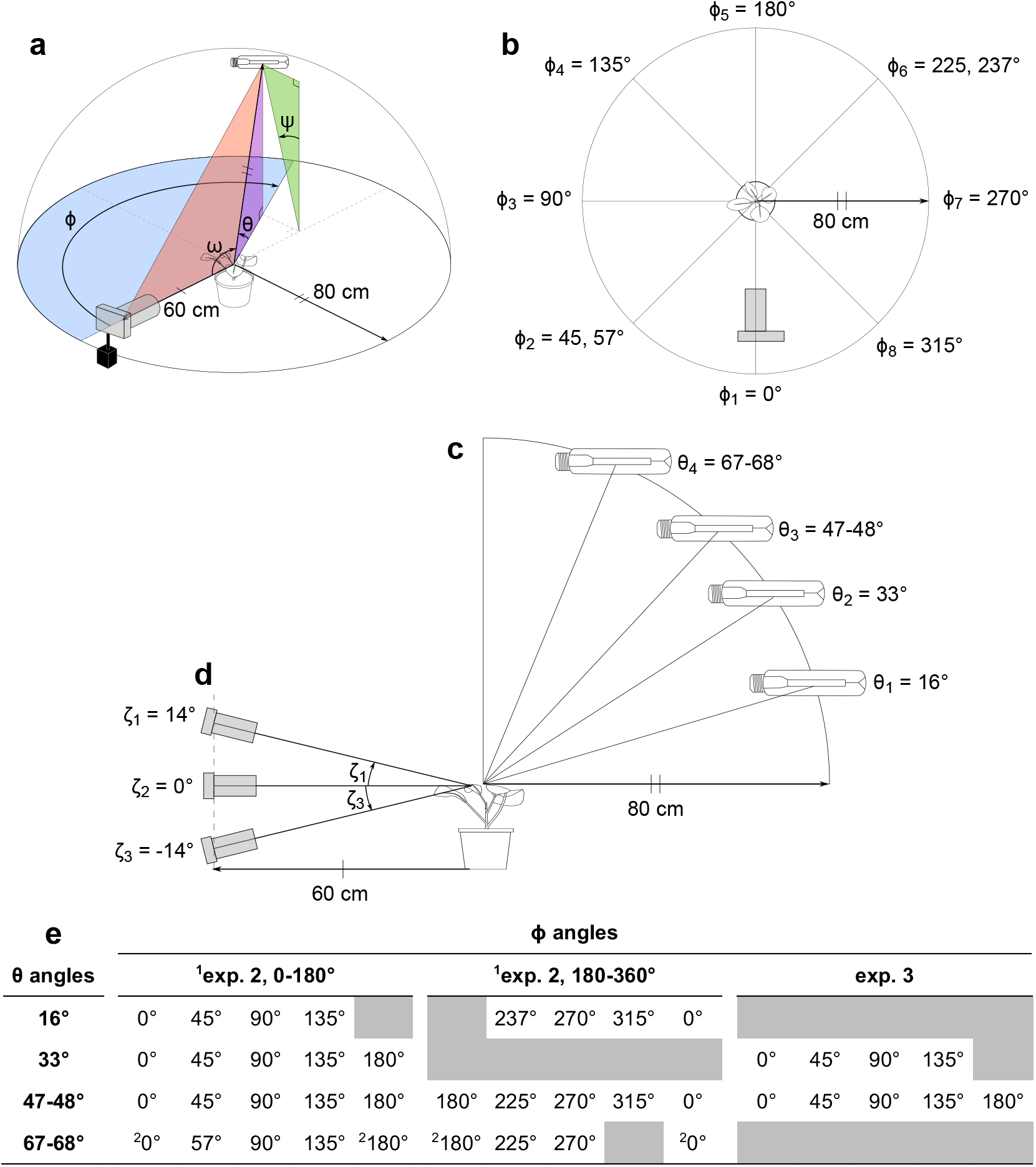
**a** Diagram showing the relative position of the camera, experimental plant and light source as well as the angles between them. The differences in azimuth between the camera and the light source (ϕ), and the elevation of the light source (θ), were manipulated to produce a range of values in the angles ω & ψ. **— b** The range of values for the angle ϕ. **— c** The range of values for the angle θ. **— d** The range of values of camera inclination (ζ). — **e** The degree of linear polarization (*DoLP*) and axis of polarization (*AoP*) were measured using photo polarimetry at each combination of ϕ and θ angles listed in the table for experiments 2 and 3. Due to restrictions of the scaffolding for mounting the metal halide lamp, certain combinations of ϕ and θ were impractical for polarimetry (shown in dark grey). For similar reasons, measurements in experiment 3 were limited to a subset of θ angles, but for each of the ϕ and θ combinations listed in the table, measurements were taken at each ζ value. ^1^In experiment 2, plants were either photographed at a ϕ between 0-180° or 180-360°. ^2^Due to low *DoLP*, these combinations were excluded from *AoP* analyses.

As *DoLP* is an important host plant cue, at least for female *P. rapae* (Blake et al., 2019), it would be informative to compare the *DoLP* and *AoP* of multiple host and non-host plants. While the polarization of select plant species has previously been examined (Grant et al., 1993), and photo polarimetry has been used to examine plant surfaces (Horváth et al., 2002), photo polarimetry has not yet been used to compare foliar reflected polarized light among different plant species. Moreover, polarization characteristics of foliage in the ultraviolet range (UV, 320-400 nm) have been predicted to resemble those in the human-visible range (400-700 nm) (Horváth et al., 2002), but this prediction has never been experimentally tested. Therefore, our first objective was to use photo polarimetry to characterize the *DoLP* and *AoP* of foliar reflections from host and non-host plants of *P. rapae* and to compare polarization characteristics of foliage in both the UV and human-visible range.

Further knowledge gaps pertain to the question as to how interspecific differences in foliar polarization are affected by the position of the observer and the light source. Positional effects have been investigated in relation to single leaves (Hegedüs & Horváth, 2004; Horváth et al., 2002) but not whole plants. Therefore, our second objective was to use photo polarimetry to measure select plant species under a series of light source and observer positions in order to model how approach trajectory affects foliar *AoP* and *DoLP*, and thus plant attractiveness, to host-plant-seeking female *P. rapae*.

## Methods

### Plant material

Within a greenhouse, we grew plants in pots (12.7 cm diam), thinning to one plant per pot except for fall rye and oregano. In these species, multiple plants per pot generated a leaf area more comparable to that of the other species examined (Table S1). Plants selected for photography in experiments were 10-20 cm tall with 4-6 fully expanded true leaves (BBCH 14-16).

### Polarimetry of Experimental Plants

We used photo polarimetry (Foster et al., 2018; Horváth & Varjú, 2004) to measure the intensity (*I*), *DoLP* and *AoP* of the selected plants. To obtain these measurements, we used a modified Olympus E-PM1 camera (Olympus, Tokyo, Japan) with expanded sensitivity in the UV (320-400 nm) (Fig S1c; Dr. Klaus Schmitt, Weinheim, Germany, uvir.eu) and an ultra-broadband linear polarizing filter (68-751, Edmund Optics, USA). We narrowed sensitivity to the human-visible range (400-700 nm) and the UV range with a UV/IR filter (Baader Plantarium, Mammendorf, Germany) and a U-filter (Baader Plantarium), respectively. To calculate the *DoLP* and the *AoP*, we took four images with the polarizing filter positioned at 0°, 45°, 90° and 135°.

We kept the white-balance, aperture, and other exposure controls constant between exposures, with all images captured in a raw image format. Within the image-analysis software platform Fiji (Schindelin et al., 2012), we used a series of custom-created macros for image analysis and measurement (Blake et al., 2020b). We decoded images with DCRAW (Coffin, 2019) as a 16-bit linear bitmap, persevering sensor linearity. We determined color corrections necessary to ensure accurate color representation through photographing a 99% Spectralon reflectance standard (SRS-99-010, Labsphere, NH, USA) under similar lighting conditions as the experimental plants (Blake et al., 2020b). We aligned all images (0°, 45°, 90°, 135°) from each plant using TurboReg (Thévenaz et al., 1998) and separated the plant in each image from the background (see below). We then calculated Stokes parameters (including *I*), *DoLP*, and *AoP* for each pixel in the red (575-700 nm), green (455-610 nm), blue (410-530 nm) and UV (330-395 nm) bands of the electromagnetic spectrum (Fig. S1c) and averaged all pixel values to give a whole-plant mean for both the intensity (*I*) and the *DoLP*, and a modal value for the *AoP*.

### Interspecific comparisons of foliar reflectance (Exp. 1)

We photographed plants upright inside a black velvet-lined box lit by a 400 W Hortilux® Blue metal halide lamp (MT400D/BUD/HTL-BLUE, EYE Lighting Int., Mentor, OH, USA) suspended 75-80 cm above the box (Fig. S2). Light was directed onto a plant by a white-cardstock tube (12.5 × 21.6 cm), thus minimizing reflections from the box walls. The camera was positioned 75-80 cm from the plant at approximately the same height as the plant canopy (Fig. S2).

In all exposures (0°, 45°, 90°, 135°) and color bands (UV, blue, green, red), we used a background mask to isolate the plant from the background. We created the background mask using areas above ~2.3% of the maximum pixel value in the green band. To eliminate possible effects of shading or unequal areas of the plants being directly lit, we limited estimations of *DoLP* and *AoP* to areas of the image above 5% of the maximum pixel value in each color band. We further limited estimates of *AoP*, in this and subsequent experiments, to areas with a *DoLP* greater than 15%, as below this *DoLP* estimates of *AoP* have little meaning (Horváth & Varjú, 1997).

### Effect of light source azimuth and elevation on foliar polarization (Exp. 2)

To photograph plants in various light source elevation and azimuth combinations (Fig. 1abce), we used scaffolding to precisely vary the height of the metal halide lamp and a movable platform to keep the camera and plant in orientation. Subtle variations in plant height did result in some variation in light source elevation but these variations and those of related angles were incorporated into the analyses. We positioned a black velvet background behind the plant in each image to enable optimal separation of the plant from the background. We took these measurements using a subset of the species we examined in the previous experiment, selecting plants with shiny leaves (potato, white mustard), matte leaves (cabbage, rutabaga) and fall rye, which holds its leaves in a more vertical orientation. We omitted UV polarimetry in this and the subsequent experiment because plants would shift position due to positive phototropism (Koller, 2000) during the extended time frame needed for several long UV exposures. Omitting UV polarimetry in experiment 2 was further justified given the strong correlation (R^2^ = 75%) between *DoLP* in the UV and blue found in experiment 1 (see Results).

As the intensity of the black velvet background varied considerably with the position of the metal halide lamp, we could not specify a single intensity threshold to separate the plant from the background as we had in the previous experiment. We therefore used a combination of all three human visual color bands to manually create a background mask. As we wanted to compare the plant in different light source positions, we estimated *DoLP* from the same subset of pixels specified by the background mask rather than limiting *DoLP* to areas with a specific intensity, as in the previous experiment.

### Effect of observer elevation on foliar polarization (Exp. 3)

Using cabbage and white mustard, we applied the same procedure as described above to examine the effect of observer elevation (camera in this case). At each observer elevation (14°, 0°, −14°), we photographed the plant at a subset of the combinations of elevation and azimuth mentioned above (Fig. 1de).

### Statistical analysis

We compared foliar reflection among species (Exp. 1), using a linear model with post-hoc Tukey’s test (Table S2; Blake et al., 2020b). We analyzed the effects of light source and observer positions (Exps. 2, 3) on foliar polarization, using mixed models with plant included as a random effect (Table S2; Bates et al., 2015). We incorporated ψ into models of *DoLP* as the square of its cosine, whereas ω was incorporated in these models via *p*(ω) as described in the Fresnel equations below (1-3), with *n*_1_ being the refractive index of air (1.00) and *n*2 being the refractive index of the leaf surface (1.34-1.79, depending on color band). For each color band, we chose the leaf surface refractive index that minimized model deviance (Blake et al., 2020b). In modeling the effect of observer elevation (Exp. 3), we incorporated ζ into existing models from Exp. 2 as its arctangent, and scaled ζ by a factor of 16 so its effect would quickly reach an asymptote as ζ moved away from 0 (Table S2; Blake et al., 2020b).

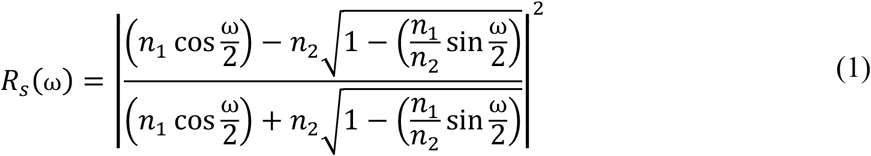

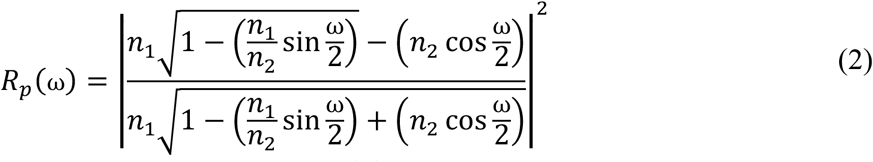

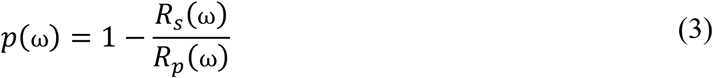

### Modeling the effect of solar elevation and azimuth on host attractiveness to *Pieris rapae*

Utilizing the models for *DoLP* and *AoP* from Exp. 3 (Table S2), we predicted *DoLP* and *AoP* across most possible values of ζ (−15–90°), all possible values of ϕ (0–360°), and a selection of θ values (15, 45, 75°; Blake et al., 2020b). These predictions were limited to the blue color channel as there were insufficient data to fit *AoP* models for the red and green color bands. Then using the ranges of *DoLP* and *AoP* shown to be unattractive to *P. rapae* (Blake et al., 2019), we modeled approach trajectories that would result in attractive and unattractive polarization characteristics, as well as low *DoLP* (<10%, moderately attractive).

## Results

### Interspecific comparisons of foliar reflectance (Exp. 1)

There were statistically significant differences in both intensity and *DoLP* among plant species in all color bands (Figs. 2ab, S3ab, S4ab, S5ab; Table S2). In contrast, we found minimal, although sometimes statistically significant, differences in *AoP* among plant species (Figs. 2c, S3c, S4c, S5c; Table S2). Differences in intensity and *DoLP* were comparably large in the UV and blue color bands. The comparatively shiny-leaved species had a much higher *DoLP* than the matt-leaved species, but only in the blue and UV bands (Figs. 2b, S5b), where most *P. rapae* host plants grouped together.

**Figure 2.**
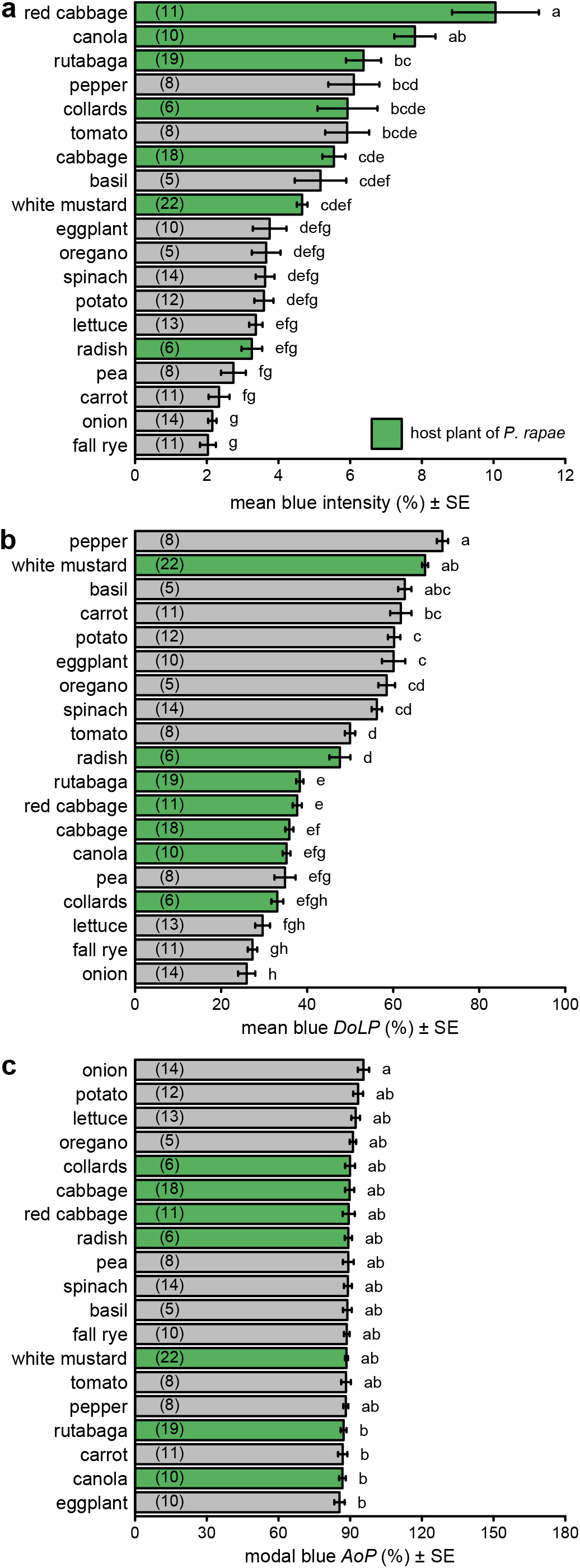
Comparison of intensity (**a**), degree of linear polarization (*DoLP*) (**b**), and axis of polarization (*AoP*) (**c**) among host plants (green bars) and non-host plants (grey bars) of *Pieris rapae*. These measurements used the blue color band, whereas measurements with other color bands are presented in Figs. S3–5. Bars show mean or modal values with the number of plants measured noted in parentheses in each bar. In each subpanel, bars with different letters differ statistically (*p*<0.05), as determined by a post-hoc Tukey test. Data in subpanels **b** and **c** were previously reported (Blake et al. 2019).

### Effect of light source azimuth and elevation on foliar polarization (Exp. 2)

For all three color bands, there was a strong relationship between ω and *DoLP* (Figs. 3, S6, S7; Table S2), with *DoLP* increasing as ω approached double the Brewster’s angle (53-60°). This relationship was less pronounced when the plants were lit more from the side (larger Ψ angle). Fall rye with mostly vertical leaf orientation showed a different and weaker relationship between ω and *DoLP* (Figs. 3a, S6a, S7a).

**Figure 3.**
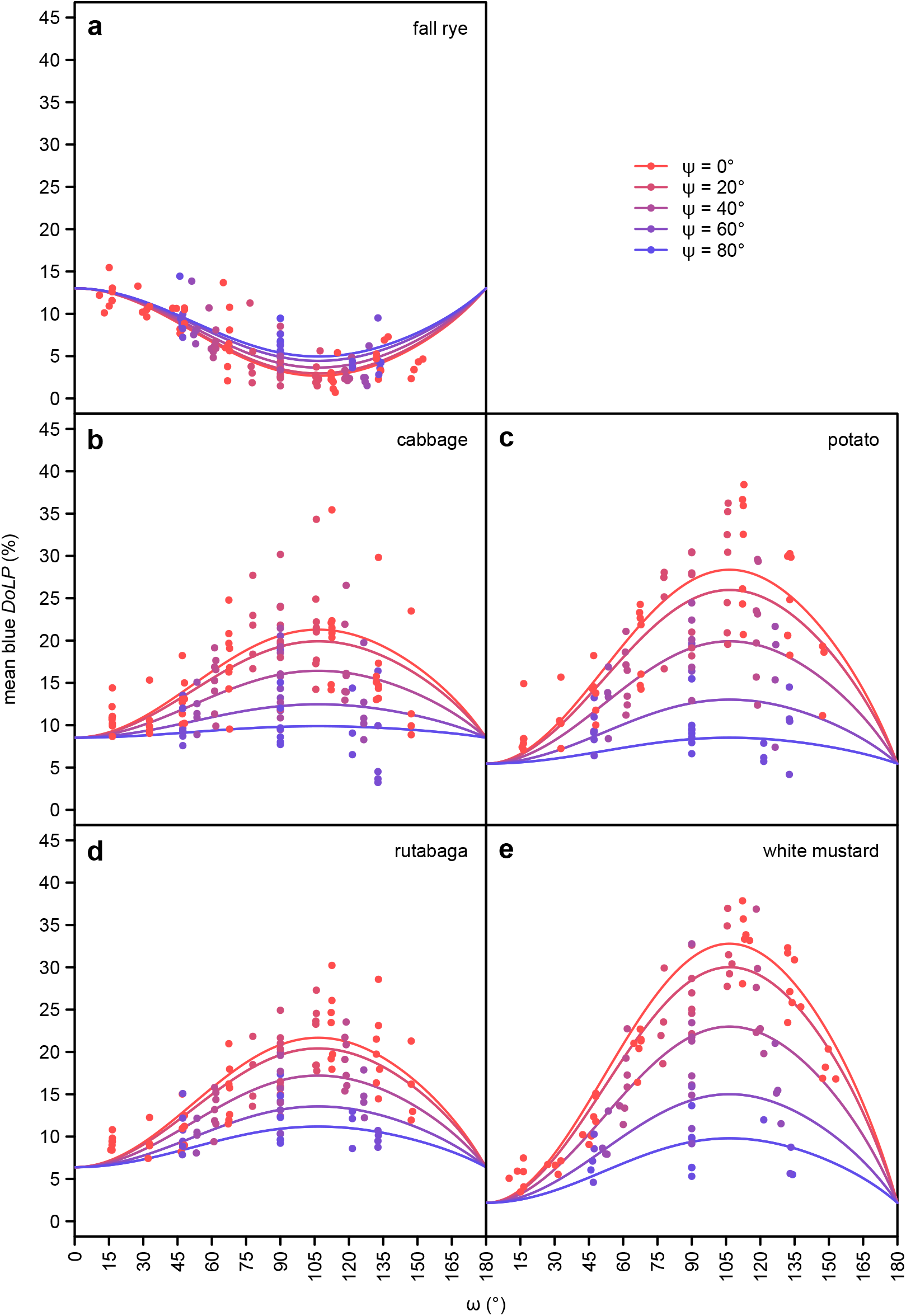
The effect of ω (angle between observer and light source with the plant at its vertex; see Fig. 1) and ψ (2-dimensional component of ω perpendicular to the plane passing through both the observer and the plant; see Fig. 1) on the mean degree of linear polarization (*DoLP*) of the blue color band, as measured in five select plant species using photo polarimetry. Data with other color bands are presented in Figs. S6–7 and show a similar relationship. Cabbage, rutabaga and white mustard are host plants of *Pieris rapae*.

There was an approximately proportional negative relationship between the ψ angle and *AoP* in all color bands (Figs. 4, S8, S9; Table S2). The slope of this relationship was steepest when the light source was behind the observer (ϕ = 0).

**Figure 4.**
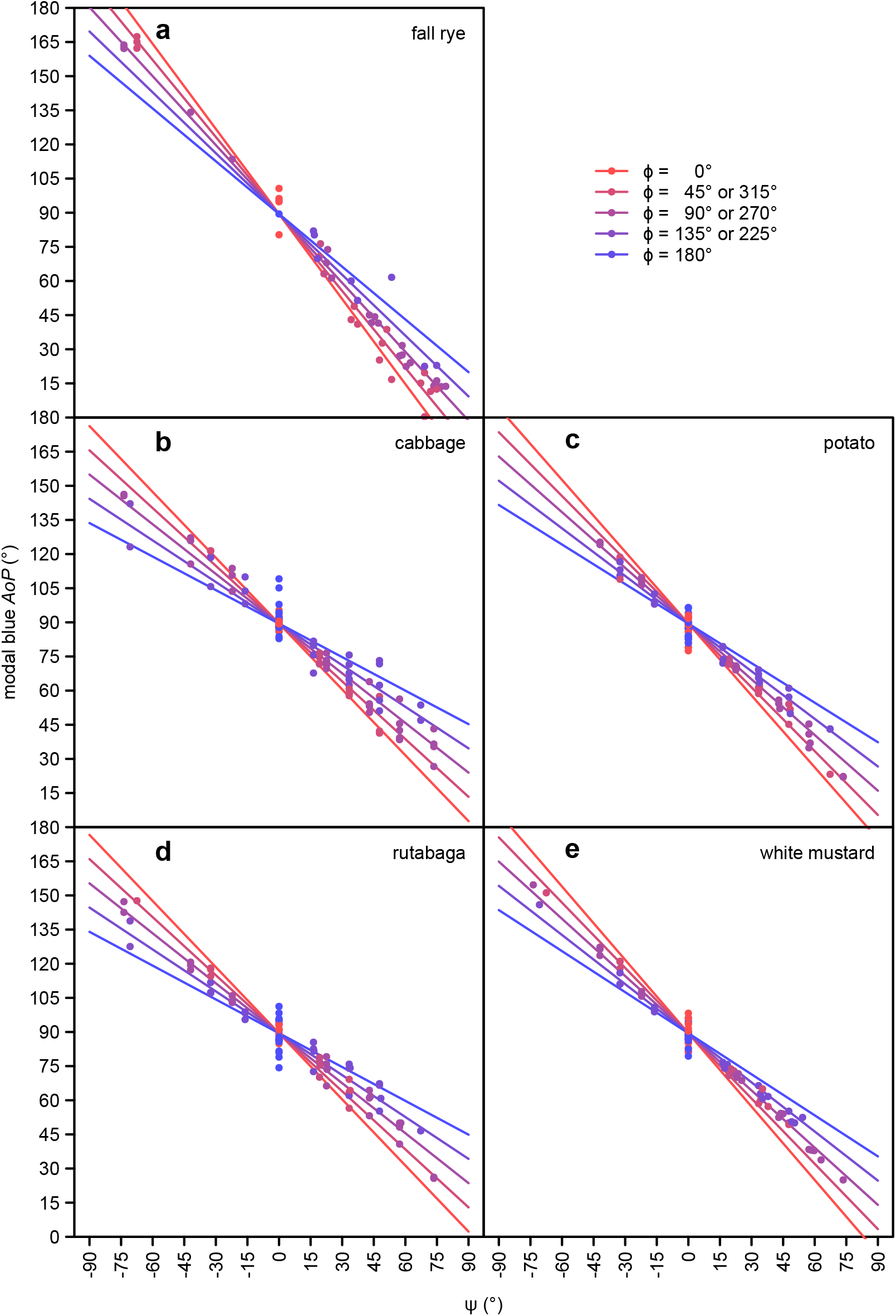
The effect of ψ (2-dimensional component of ω perpendicular to the plane passing through both the observer and the plant; see Fig. 1) and ϕ (angle between the azimuth of the observer and the light source; see Fig. 1) on the modal axis of polarization (*AoP*) of the blue color band, as measured in five select plant species using photo polarimetry. Data of other color bands are presented in Figs. S8–9 and show a similar relationship. Cabbage, rutabaga and white mustard are host plants of *Pieris rapae*.

### Effect of observer elevation on foliar polarization (Exp. 3)

The effect of ω on *DoLP* increased with the elevation of the observer (ζ; Figs. 5, S10, S11; Table S2). The elevation of the observer also affected *AoP* (Fig. S12). The slope of the relationship between the ψ angle and *AoP* was shallower at lower observer elevations, while the effect of the ϕ angle on the relationship been ψ angle and *AoP* was more pronounced at higher observer elevations. These effects were all relatively subtle in comparison to the effects of light source position.

**Figure 5.**
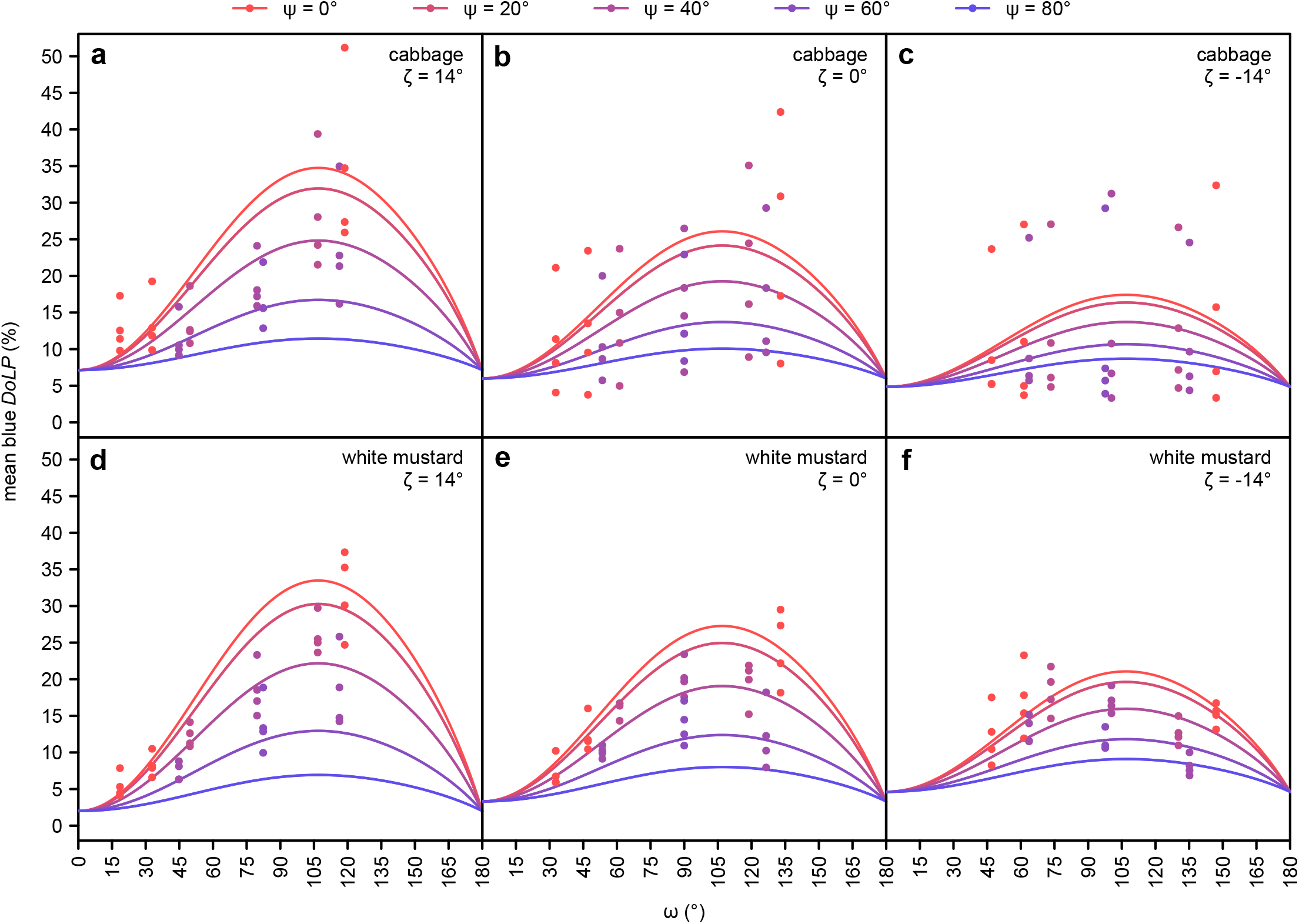
The additional effect of observer elevation (ζ; see Fig. 1) on the mean degree of linear polarization (*DoLP*) of the blue color band, as measured in cabbage and white mustard (host plants of *Pieris rapae*) using photo polarimetry. Data with other color bands are presented in Figs. S10–11 and show a similar relationship.

### Modeling the effect of solar elevation and azimuth on host plant attractiveness to *Pieris rapae*

As indicated by our modeling, the greatest *DoLP* of foliage is realized when the light source is located directly behind the plant (Figs 6a-c, S13a-c). Effects of solar elevation (θ) on *DoLP* could be compensated for, in part, by shifting the observer elevation (ζ) but lower observer elevation reduced overall *DoLP*.

**Figure 6.**
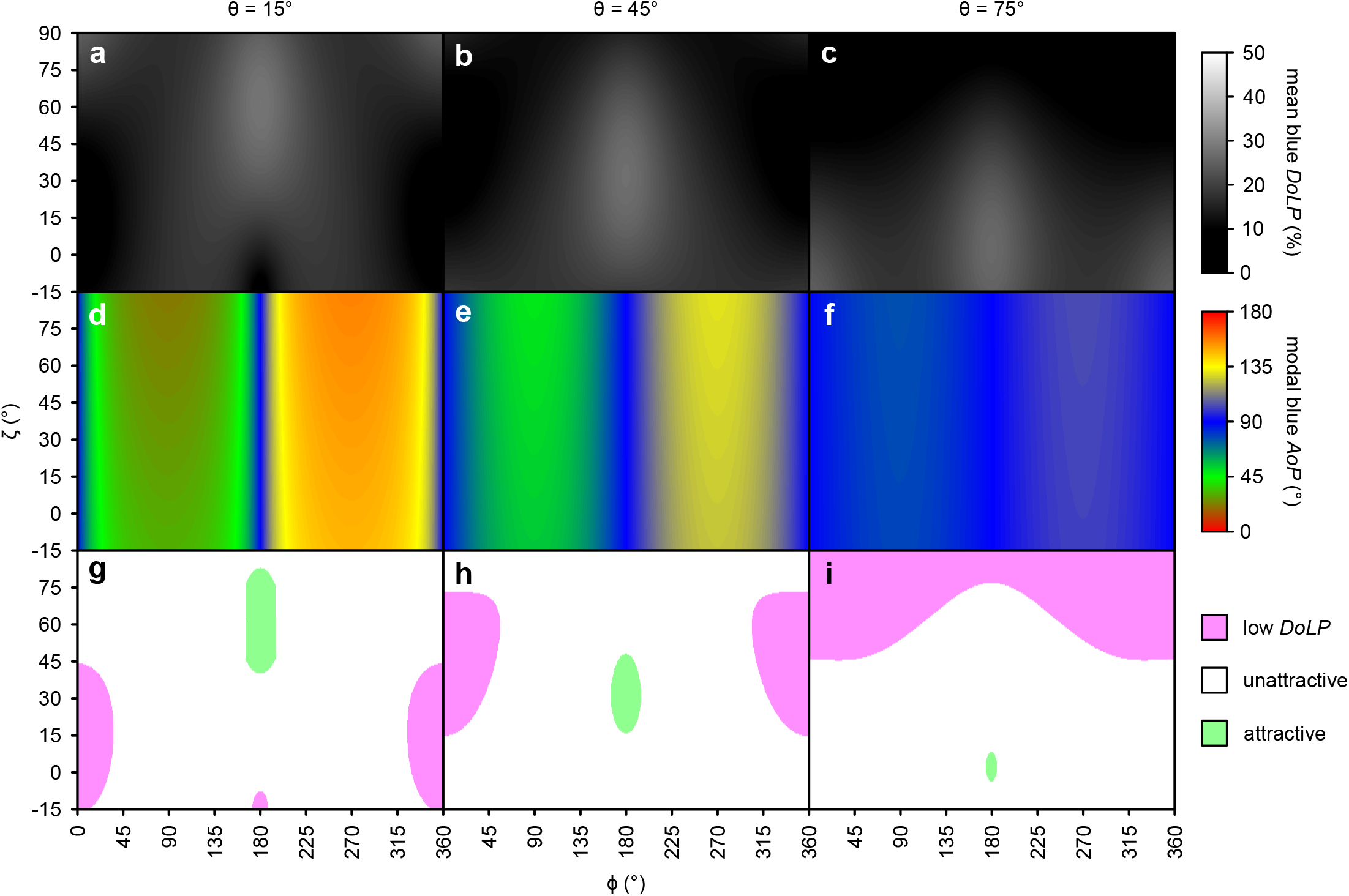
Effects of approach direction (angle between the azimuth of the observer and the light source (ϕ; see Fig. 1) and elevation of the observer (ζ; see Fig. 1) on the mean degree of linear polarization (*DoLP*) (**a-c**) and the modal axis of polarization (*AoP*) (**d-f**) of the blue color band of cabbage plants (host of *Pieris rapae*). Attractiveness of resulting polarization characteristics to *P. rapae* (**g-i**), based on a previous behavioral study (Blake et al. 2019). Approach trajectories resulting in attractive characteristics (*DoLP* = 26-36% and *AoP* = 0-38, 53-128 or 143-180°) and unattractive characteristics (*DoLP* = 10-26% or *AoP* = 38-53°, 128-143°) are shown in green and white, respectively, with pink indicating trajectories resulting in a moderately-attractive low *DoLP* (<10%). Higher *DoLP* (36-60%) would also be unattractive but were not predicted by these models. These effects changed with light source elevation (θ; see Fig. 1) which is shown at 15° (**a, d, g**), 45° (**b, e, h**) and 75° (**c, f, i**). Analogous data were obtained with white mustard (Fig. S13), another host plant of *Pieris rapae*.

Our model predicts that the greatest range of *AoP* across all ϕ angles tested is found when solar elevation (θ) is low, with ϕ angles at or near 180° always yielding an *AoP* near 90°. We also note that smaller shifts in *AoP* occur with ϕ angle at lower observer angles (ζ), but this effect is relatively small.

When we modeled ζ, ϕ and θ values resulting in combinations of *DoLP* and *AoP* attractive and unattractive to *P. rapae* (Figs. 6, S13), there was consistently a window of attractive *DoLPs* at a ϕ angle of 180°, and a moderately attractive low *DoLP* area opposite it at a ϕ angle of 0°. All other combinations of ϕ and ζ resulted in unattractive *DoLPs*. Increasing solar elevation (θ) shifted the attractive window downward and the low *DoLP* area upwards. Increased solar elevation (θ) also decreased the size of the attractive window, while increasing the size of the low *DoLP* area. The *AoP* had little effect on these windows outside of a small narrowing of the attractive window at low solar elevations (θ).

## Discussion

Our study confirms earlier work demonstrating large differences in *DoLP* among plant species (Blake et al., 2019; Grant et al., 1993) and refines our understanding of polarized reflections from plant foliage. Unlike previous studies that examined polarized reflections of single leaves, models of leaves, or plant canopies (Grant et al., 1993; Hegedüs & Horváth, 2004; Horváth et al., 2002; Horváth & Hegedüs, 2014; Maignan et al., 2009; Raven, 2002; Rondeaux & Herman, 1991; Vanderbilt & Grant, 1985; Woolley, 1971), we recorded reflections from entire plants thereby revealing several emergent phenomena. Most importantly, our modeling suggests that certain approach trajectories are optimal for foraging insects to discriminate among plant species based on the *DoLP* of foliar reflections.

Our measurements of polarization of foliar reflections are consistent with point-source polarimetry data (Grant et al., 1993), and other photo polarimetry of plant surfaces (Fig. 2, S3–5; Hegedüs and Horváth, 2004; Horváth et al., 2002). As predicted by Horváth et al. (2002), our UV polarimetry data closely resemble those of the human-visible color bands, especially blue, and are consistent with previous measurements in the human-visible range. Similar to previous measurements (Grant et al., 1993; Horváth et al., 2002), glossy, flat and/or dark leaf surfaces have an increased ratio of specular to diffuse reflection and greater *DoLP* than matte, undulating, and/or bright leaf surfaces. As leaves have low reflectance in the green and red color bands, the *DoLP*s in the blue and UV color bands expectedly exceeded those in the green and red bands. Moreover, plants with leaves that tend to be held more vertically (e.g., fall rye, onions), and provide little horizonal surface for specular reflections, had low *DoLP* values. Despite large differences in leaf shape (simple *vs* compound) and growth form (all basal leaves *vs* basal and cauline leaves), there were only small, albeit statistically significant, differences in *AoP* between plant species (Table S2). These findings in combination with the smaller interspecific differences in intensity relative to *DoLP* (Table S2), further support the conclusion that foliar *DoLP* is *the* visual cue that conveys the most host plant information, especially information about the foliar surface (waxes, pubescence, undulations).

The angle between light source, plant and observer (ω) strongly predicted the foliar *DoLP* (Figs. 3, S6, S7) for all color bands, with the strongest polarization at twice the Brewster’s angle (53-60°). These data are consistent with both theoretical predictions and other experimental measurements of the effect of viewing angles on *DoLP* (Horváth et al., 2002; Raven, 2002; Rondeaux & Herman, 1991; Woolley, 1971). However, the phenomenon of lowering the *DoLP* with increasing Ψ had not previously been noted and emerges here through whole-plant measurements incorporating multiple leaf surfaces. As the orientation of plant leaves is typically more horizontal than vertical, but not perfectly horizontal, plants lit more from the side than from above (greater ψ) have a relatively greater leaf area shadowed by their own leaves. These shadowed areas have a lower *DoLP*, lowering the plants’ overall *DoLP*. Of course, this relationship was absent in fall rye (at least at the growth stage examined) with primarily vertically held leaves. When plants were photographed at or below the level of the leaf canopy (lower ζ), the *DoLP* was reduced (Figs. 5, S10, S11). Similar to the effect of ψ, lower ζ results in a smaller leaf surface reflecting light at the observer, and a larger leaf surface being in shadow or showing light transmitted through the leaves. Light transmitted through leaves has a low *DoLP* due to diffuse scattering by plant tissue, as previously observed in single leaf measurements (Horváth et al., 2002; Vanderbilt & Grant, 1985).

In agreement with prior examinations of foliar polarization, the *AoP* of all color bands was largely a function of ψ (Figs. 4, S8, S9), with values of *AoP* moving away from 90° as the light source was less aligned with the line between the observer and the plant (see Fig. 1a). This relation between *AoP* and ψ is consistent with previous observations (Horváth et al., 2002; Können, 1985). Although the *AoP* of a particular plant area did not change much in relation to the light source position, the variety of leaf orientations within a single plant and the curvature of leaf surfaces ensured that at least a portion of the plant showed a specular reflection regardless of the light source’s position relative to the plant. Invariably, these areas of specular reflection showed a greater *DoLP* accounting for much of the observed relationship between *AoP* and ψ. The variety of leaf surface orientations and the resultant *AoPs* also explains why the relationship between *AoP* and ψ is shallower than the inversely proportional relationship one could expect. When plants were viewed with the light source directly in front of the observer (ϕ = 180°; Figs. 4, S8, S9), the relationship between *AoP* and ψ had a reduced slope, a phenomenon being more pronounced when the plant was observed from a higher angle (ζ > 0; Fig. S12). In both cases (ϕ = 180°, ζ > 0), this resulted in plants having a higher overall *DoLP* (Figs. 3, 5), and consequently less leaf surface area (with a < 15% *DoLP*) being excluded from estimations of *AoP*. Given that less polarized leaf surface areas showed a weaker relationship between *AoP* and ψ, the overall lower *DoLP* resulted in a stronger relationship between *AoP* and ψ as only leaf areas with highest *DoLP* were above the cutoff. All these effects of ψ on *AoP* could potentially have biological relevance if a host plant foraging insect were to weigh observations of *AoP* by their *DoLP* when determining a plant’s overall *AoP*. Nonetheless, in our modeling, these specific effects on *AoP*, and the effects of *AoP* in general, on host plant attractiveness to *P. rapae* seem to be subtle in comparison to the effects of *DoLP* (Figs. 6, S13).

Our modeling of the effect of approach trajectory on visual attractiveness of plants to *P. rapae* revealed that *DoLP* is a much more important determinant of plant attractiveness than *AoP* (Fig. 6). Approach trajectories resulting in *AoP* unattractive to *P. rapae* were also unattractive due to low *DoLP*. It follows that the effect of *AoP* on plant attractiveness can largely be discounted. The key determinant of an attractive *DoLP* was the azimuth of an approach trajectory relative to the light source (ϕ). This was due to its effect on ψ, as plants obliquely lit even at the Brewster’s angle showed a much lower *DoLP*. In fact, the only attractive approach trajectories were those where the light source was located behind the target plant. *DoLP* and attractiveness were also affected by how close the angle between observer, plant and light source (ω) was to twice the Brewster’s angle, which is affected by light source elevation (θ), observer elevation (ζ), and azimuth (ϕ). However, when the light source was behind the plant, there was always a combination of θ and ζ allowing for foliar reflections approaching the Brewster’s angle. Although high solar elevations (>75°) – constrained to times near solar noon and limited to latitudes near the equator – are relatively rare, they would require much lower approach angles for accurate assessment of foliar *DoLP*. It is the key result of our modelling that for most solar positions there is a single optimal approach trajectory that would best enable a foraging insect to assess foliar *DoLP*. However, this conclusion applies only to settings where foliar reflections are dominated by the specular reflections of sunlight (or another single strong unpolarized light source), as we took measurements indoors and did not incorporate possible effects of polarized skylight (Hegedüs & Horváth, 2004; Horváth et al., 2002).

We have every reason to predict that our polarization modeling is applicable to the foliar reflectance of many plant species. However, the data we have obtained with herbaceous flowering plants may not be applicable to graminoids, such as fall rye, or other plants with more vertically held leaves. Moreover, due to the size of trees and large shrubs, foraging insects more often approach them from below (reference), and do not view them in their entirety, complicating the applicability of our modeling. It would therefore be intriguing to model whether approach trajectories have similar effects on polarized light cues that may be used by insect herbivores of trees and shrubs.

While this work focused on *P. rapae*, our *DoLP* and *AoP* modeling should be applicable to other polarization-sensitive visual systems. Furthermore, our prediction of a single optimal approach trajectory for the discrimination of *DoLP* should hold true for other polarization-sensitive insects such as *Papilio* butterflies (Kelber et al., 2001; Kinoshita et al., 2011), where increased *DoLP* of foliar reflections would be expected to have a linear effect on attractiveness (Blake et al., 2020a). The approach of butterflies to host plants has not yet been well documented and – accordingly – no stereotyped approach has been noted, as one would anticipate based on our predictions of polarized reflections. Reminiscent of the plunge responses of *Notonecta* backswimmers (Schwind, 1984), one might expect an approach where the butterflies’ trajectory is constrained so that at least a portion of the compound eyes are viewing the plant at or near the Brewster’s angle. Alternatively, butterflies might circle plants before landing, thereby shifting their azimuth relative to sun, and entering and exiting the attractive window we identified. Circling plants would also allow for sequential comparison of visual information from the plant surface, aiding in *DoLP* assessment through differences in color and/or intensity (Horváth & Varjú, 2004). Mapping the position of butterflies in a 3-dimensional space during approaches to host plants would give insight into how these insects perceive and use the plants’ polarized light cues.

In conclusion, using photo polarimetry to examine polarized reflections from entire plants, we show that host and non-host plants of *P. rapae* differ in the *DoLP* of foliar reflections, with UV measurements closely resembling those of blue. Our photo polarimetry further reveals that there is a single optimal approach trajectory that would enable a foraging insect (or other observers) to best discriminate among these interspecific differences in polarization. This optimal approach trajectory is always in the direction of the light source but its inclination is dependent upon the elevation of the light source (θ). It would now be intriguing to determine whether the trajectories of polarization-sensitive insects towards host plants match those predicted by our models.

## Acknowledgments

This study was supported by an Alexander Graham Bell Canadian Graduate Scholarship to AJB, NSERC Undergraduate Student Research Awards to MCG and SC, and by a NSERC-Industrial Research Chair to GG, with Scotts Canada Ltd. and BASF Canada Ltd. as the industrial partners.

## Data Accessibility

Data are available from Mendeley Data: http://doi.org/10.17632/5bh5mhmvrk.1 (Blake et al., 2020b).

## Author Contributions

AJB, SC, MCG performed the polarimetry; AJB designed experiments; AJB performed modeling and analyses; GG supervised the project; AJB wrote the first draft of the paper and GG provided comments.

## List of symbols and abbreviations

*I*: intensity
*DoLP*: degree of linear polarization
*AoP*: axis of polarization
R: Red (575-700 nm, see Fig. S1c)
G: Green (450-625 nm, see Fig. S1c)
B: Blue (400-525 nm, see Fig. S1c)
UV: Ultraviolet (325-400 nm, see Fig. S1c)
ϕ: angle between the azimuth of the observer and the light source (see Fig. 1)
θ: elevation of light source (see Fig. 1)
ω: angle between observer and light source with the plant at its vertex (see Fig. 1)
Ψ: 2-dimensional component of ω perpendicular to the plane passing through both the observer and plant (see Fig. 1)
ζ: elevation of the observer (see Fig. 1)

---

**Table S1.**
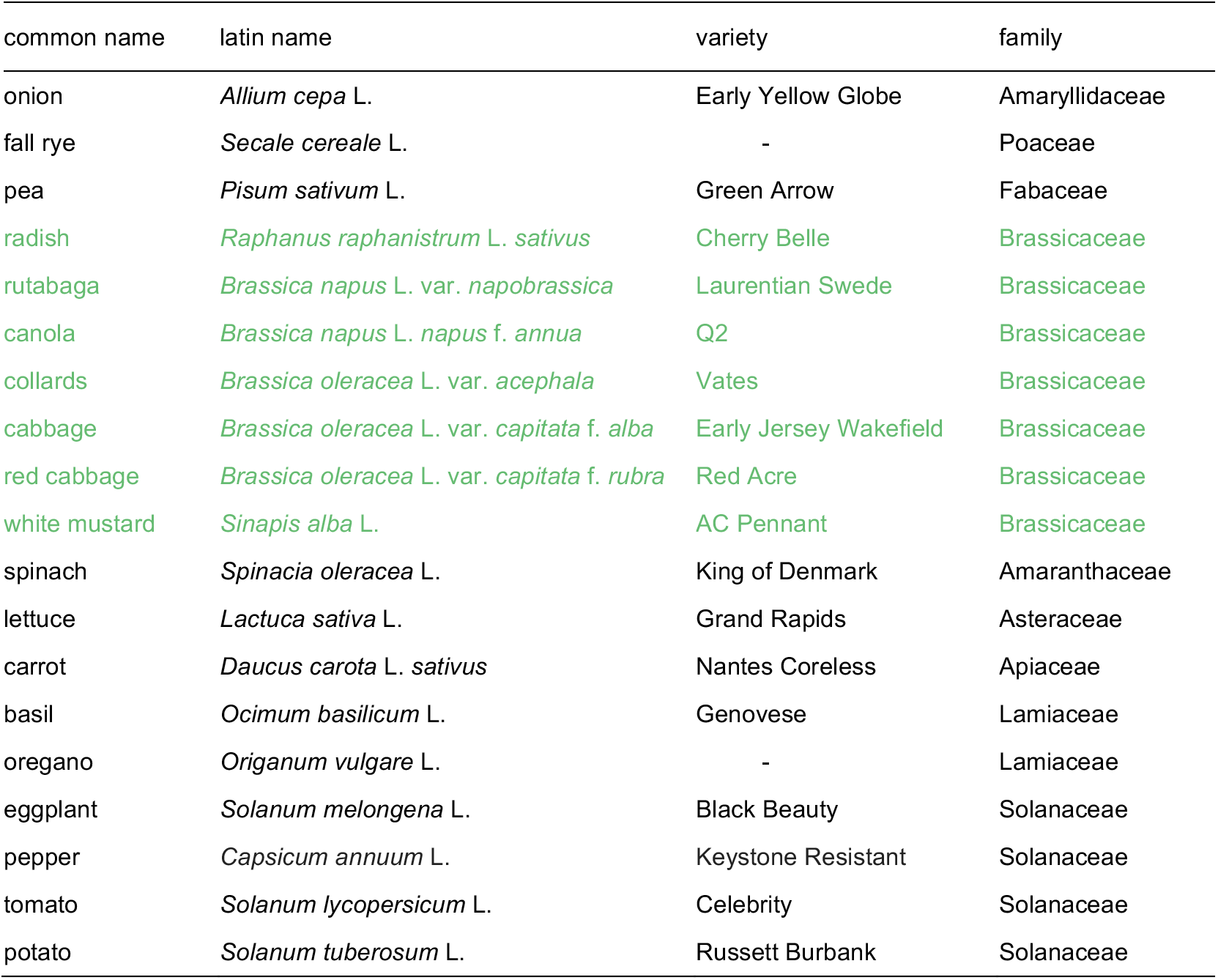
Variety and taxonomic information of select host plants (green) and non-host plants (black) of *Pieris rapae*.

| common name | latin name | variety | family |
| --- | --- | --- | --- |
| onion | *Allium cepa* L. | Early Yellow Globe | Amaryllidaceae |
| fall rye | *Secale cereale* L. | - | Poaceae |
| pea | *Pisum sativum* L. | Green Arrow | Fabaceae |
| radish | *Raphanus raphanistrum* L. *sativus* | Cherry Belle | Brassicaceae |
| rutabaga | *Brassica napus* L. var. *napobrassica* | Laurentian Swede | Brassicaceae |
| canola | *Brassica napus* L. *napus* f. *annua* | Q2 | Brassicaceae |
| collards | *Brassica oleracea* L. var. *acephala* | Vates | Brassicaceae |
| cabbage | *Brassica oleracea* L. var. *capitata* f. *alba* | Early Jersey Wakefield | Brassicaceae |
| red cabbage | *Brassica oleracea* L. var. *capitata* f. *rubra* | Red Acre | Brassicaceae |
| white mustard | *Sinapis alba* L. | AC Pennant | Brassicaceae |
| spinach | *Spinacia oleracea* L. | King of Denmark | Amaranthaceae |
| lettuce | *Lactuca sativa* L. | Grand Rapids | Asteraceae |
| carrot | *Daucus carota* L. *sativus* | Nantes Coreless | Apiaceae |
| basil | *Ocimum basilicum* L. | Genovese | Lamiaceae |
| oregano | *Origanum vulgare* L. | - | Lamiaceae |
| eggplant | *Solanum melongena* L. | Black Beauty | Solanaceae |
| pepper | *Capsicum annuum* L. | Keystone Resistant | Solanaceae |
| tomato | *Solanum lycopersicum* L. | Celebrity | Solanaceae |
| potato | *Solanum tuberosum* L. | Russett Burbank | Solanaceae |

**Table S2.** Model statements, test statistics, and p-values for statistical models of photo polarimetry determined measurements of intensity (*I*), degree of linear polarization (*DoLP*), and axis of polarization (*AoP*) for the red (R), green (G), blue (B), ultraviolet (UV, Exp. 1 only) color bands in experiments 1-3. The angles (ϕ, θ, ω, ψ, ζ) in the model statements are described in Fig. 1. The *p*(ω) relationship is defined in equations 1–3. The fixed effect of different plant species in the model is represented by *species*, whereas the random effect of individual plants was fit as an intercept and is represented by (1 | plant). The full R code used for statistical analysis is presented in an associated Dryad dataset (Blake et al., 2020b).

| experiment 1 |  |  |  |  |
| --- | --- | --- | --- | --- |
| model statement | *F* | df | *P* value |  |
| *I<sub>R</sub>* ~ *species* | 14.95 | 18,192 | < 0.0001 |  |
| *I<sub>G</sub>* ~ *species* | 15.67 | 18,192 | < 0.0001 |  |
| *I<sub>B</sub>* ~ *species* | 19.35 | 18,192 | < 0.0001 |  |
| *I<sub>UV</sub>* ~ *species* | 16.34 | 18,192 | < 0.0001 |  |
| *DoLP<sub>R</sub>* ~ *species* | 21.87 | 18,191 | < 0.0001 |  |
| *DoLP<sub>G</sub>* ~ *species* | 21.43 | 18,192 | < 0.0001 |  |
| *DoLP<sub>B</sub>* ~ *species* | 97.93 | 18,192 | < 0.0001 |  |
| *DoLP<sub>UV</sub>* ~ *species* | 72.79 | 18,181 | < 0.0001 |  |
| *AoP<sub>R</sub>* ~ *species* | 1.76 | 18,186 | 0.0334 |  |
| *AoP<sub>G</sub>* ~ *species* | 0.83 | 18,188 | 0.6625 |  |
| *AoP<sub>B</sub>* ~ *species* | 1.89 | 18,191 | 0.0186 |  |
| *AoP<sub>UV</sub>* ~ *species* | 1.02 | 18,176 | 0.4454 |  |
| experiment 2 |  |  |  |  |
| model statement | $\chi^2$ | df | *P* value | |
| *DoLP<sub>R</sub>* ~ *p*($\omega$) + *species* + *p*($\omega$) : *species* + *p*($\omega$) : $\cos \psi^2$ + *p*($\omega$) : *species* : $\cos \psi^2$ + (1 \| *plant*) | 1049 | 14 | < 0.0001 | |
| *DoLP<sub>G</sub>* ~ *p*($\omega$) + *species* + *p*($\omega$) : *species* + *p*($\omega$) : $\cos \psi^2$ + *p*($\omega$) : *species* : $\cos \psi^2$ + (1 \| *plant*) | 1059 | 14 | < 0.0001 | |
| *DoLP<sub>B</sub>* ~ *p*($\omega$) + *species* + *p*($\omega$) : *species* + *p*($\omega$) : $\cos \psi^2$ + *p*($\omega$) : *species* : $\cos \psi^2$ + (1 \| *plant*) | 1073 | 14 | < 0.0001 | |
| *AoP<sub>R</sub>* ~ $\psi$ + $\psi$ : $\phi$ + $\psi$ : *species* + (1 \| *plant*) | 912 | 5 | < 0.0001 | |
| *AoP<sub>G</sub>* ~ $\psi$ + $\psi$ : $\phi$ + $\psi$ : *species* + (1 \| *plant*) | 801 | 5 | < 0.0001 | |
| *AoP<sub>B</sub>* ~ $\psi$ + $\psi$ : $\phi$ + $\psi$ : *species* + (1 \| *plant*) | 1267 | 6 | < 0.0001 | |
| experiment 3 |  |  |  |  |
| model statement | $\chi^2$ | df | *P* value | |
| *DoLP<sub>R</sub>* ~ *p*($\omega$) + *species* + *atan*(16 · $\zeta$) + *p*($\omega$) * *species* + *p*($\omega$) : $\cos \psi^2$ + *species* : *atan*(16 · $\zeta$) + *p*($\omega$) : *species* : $\cos \psi^2$ + *p*($\omega$) : *atan*(16 · $\zeta$) : $\cos \psi^2$ + (1 \| *plant*) | 342 | 8 | < 0.0001 | |
| *DoLP<sub>G</sub>* ~ *p*($\omega$) + *species* + *atan*(16 · $\zeta$) + *p*($\omega$) * *species* + *p*($\omega$) : $\cos \psi^2$ + *species* : *atan*(16 · $\zeta$) + *p*($\omega$) : *species* : $\cos \psi^2$ + *p*($\omega$) : *atan*(16 · $\zeta$) : $\cos \psi^2$ + (1 \| *plant*) | 386 | 8 | < 0.0001 | |
| *DoLP<sub>B</sub>* ~ *p*($\omega$) + *species* + *atan*(16 · $\zeta$) + *p*($\omega$) * *species* + *p*($\omega$) : $\cos \psi^2$ + *species* : *atan*(16 · $\zeta$) + *p*($\omega$) : *species* : $\cos \psi^2$ + *p*($\omega$) : *atan*(16 · $\zeta$) : $\cos \psi^2$ + (1 \| *plant*) | 284 | 8 | < 0.0001 | |
| *AoP<sub>R</sub>* ~ $\psi$ + $\psi$ : $\phi$ + $\psi$ : *atan*(16 · $\zeta$) + $\psi$ : *species* + $\psi$ : $\phi$ : *atan*(16 · $\zeta$) + $\psi$ : *species* : *atan*(16 · $\zeta$) + (1 \| *plant*) | 419 | 6 | < 0.0001 | |

**Figure S1.**
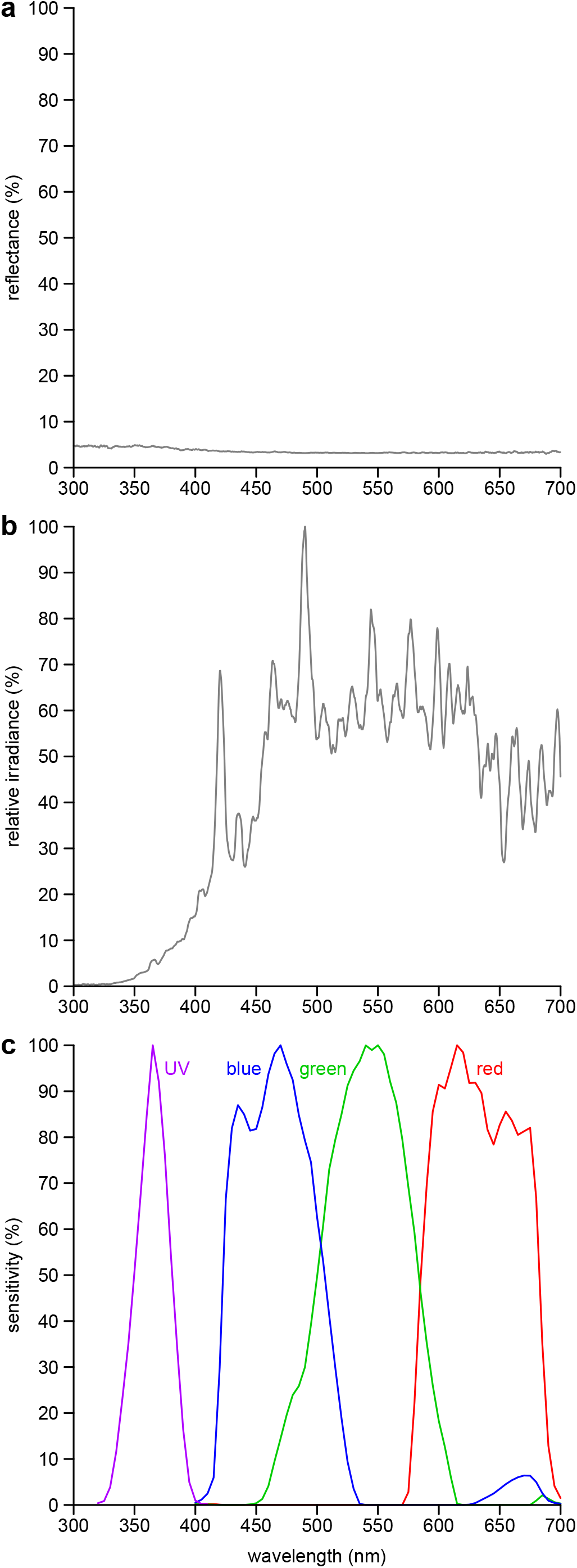
Spectra of background, illumination sources, and camera sensitivity. **a**, Reflection spectrum of the black velvet background. **b**, Relative irradiance of the metal halide lamp. **c**, Spectral sensitivity of the modified Olympus E-PM1 camera in the ultraviolet (UV), blue, green and red bands of the electromagnetic spectrum. Reflectance spectra were measured with a JAZ spectrometer (Ocean Optics Inc., Dunedin, FL, USA) calibrated with a 99% Spectralon reflectance standard (SRS-99-010, Labsphere, NH, USA). Irradiance spectra were measured with a calibrated HR-4000 spectrophotometer (Ocean Optics Inc.). Isoquantal monochromatic light for spectral sensitivity determination was generated with the same HR-4000 spectrophotometer and a scanning monochromator (MonoScan 2000, Mikropak GmbH, Ostfildern, Germany).

**Figure S2.**
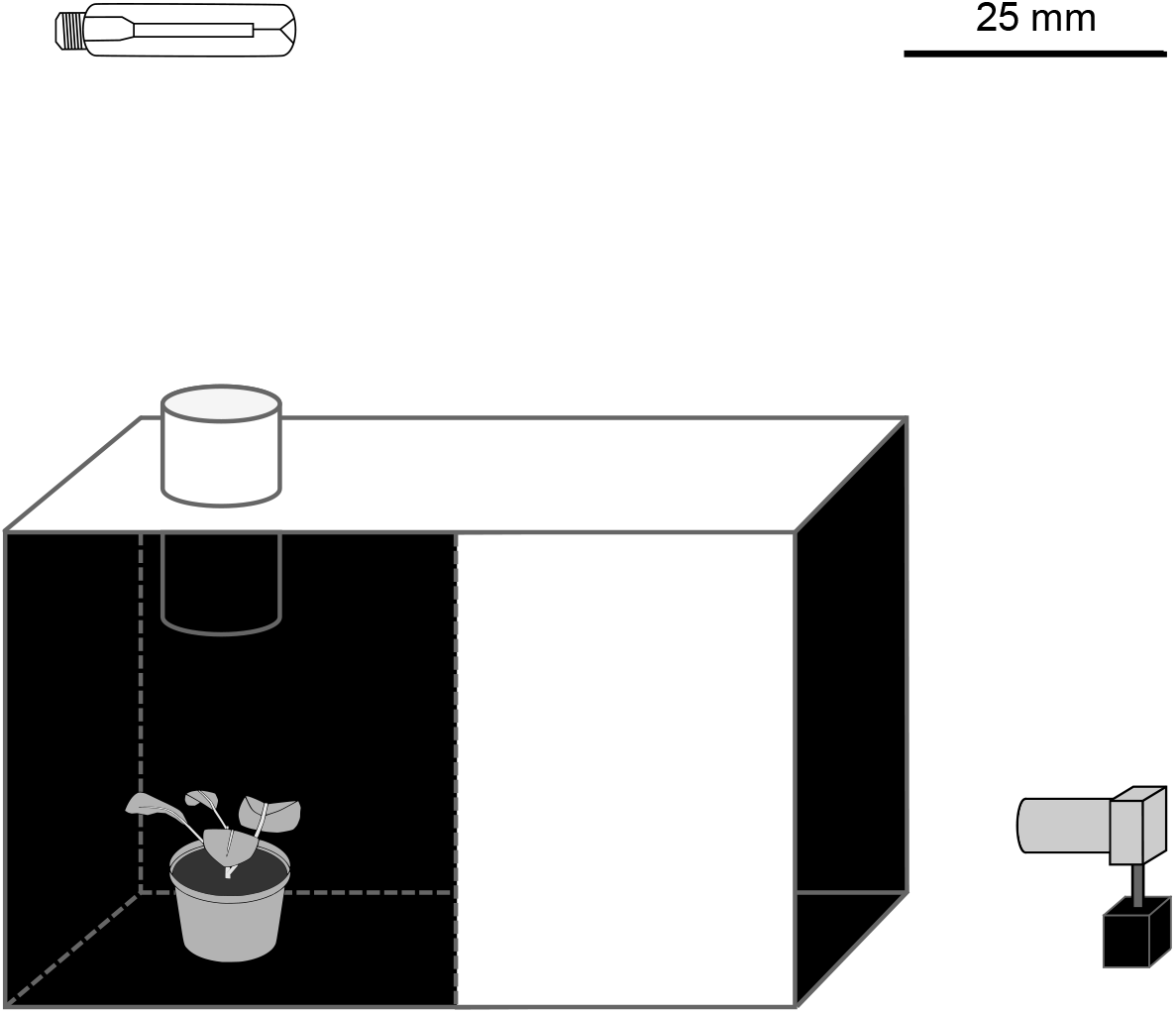
Design for photo polarimetry deployed to characterize the intensity, degree, and axis of linear polarization of various host and non-host plants of *Pieris rapae* in the red, green, blue, and ultraviolet color bands. The camera was positioned so that its optical axis was level with the plant canopy. The plant was positioned underneath the spotlight to avoid illumination of box walls. The angle between the camera and the light source was approximately 90°.

**Figure S3.**
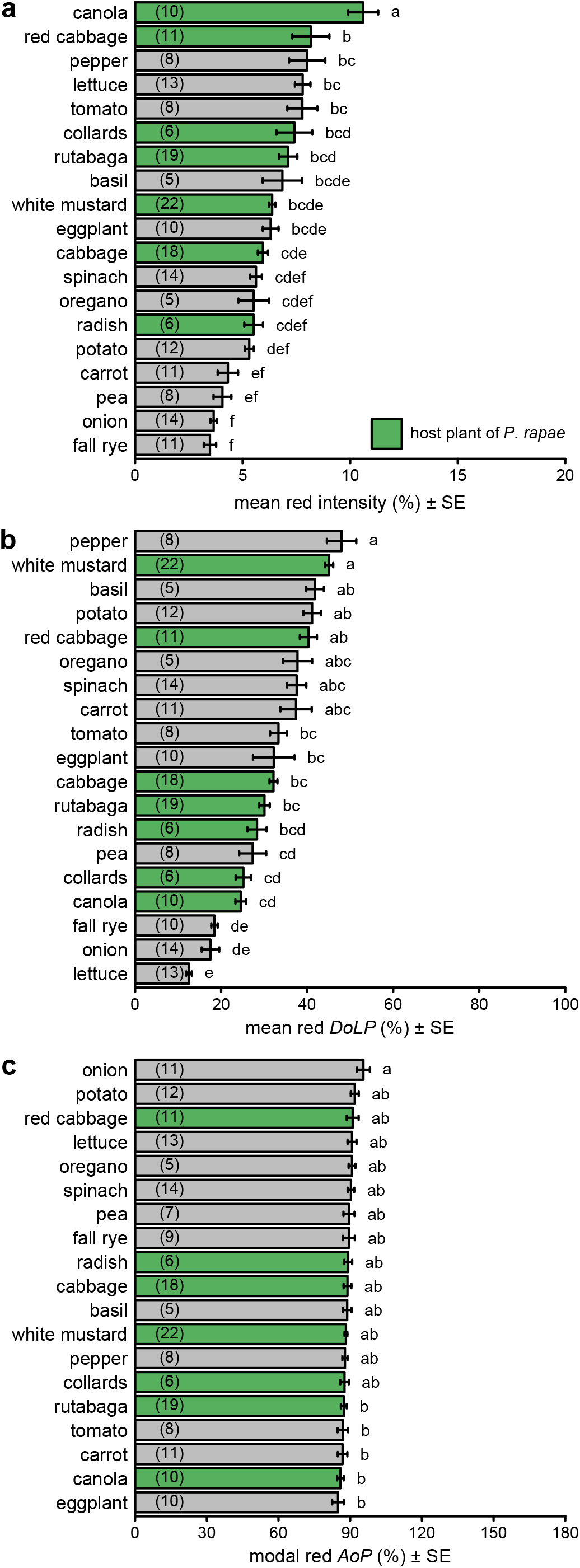
Comparison of intensity (**a**), degree of linear polarization (*DoLP*) (**b**), and axis of polarization (*AoP*) (**c**) among host plants (green bars) and non-host plants (grey bars) of *Pieris rapae*. These measurements used the red color band. Bars show mean or modal values with number of plants measured noted in parentheses in each bar. Bars with different letters differ statistically (*p*<0.05), as determined by a post-hoc Tukey test.

**Figure S4.**
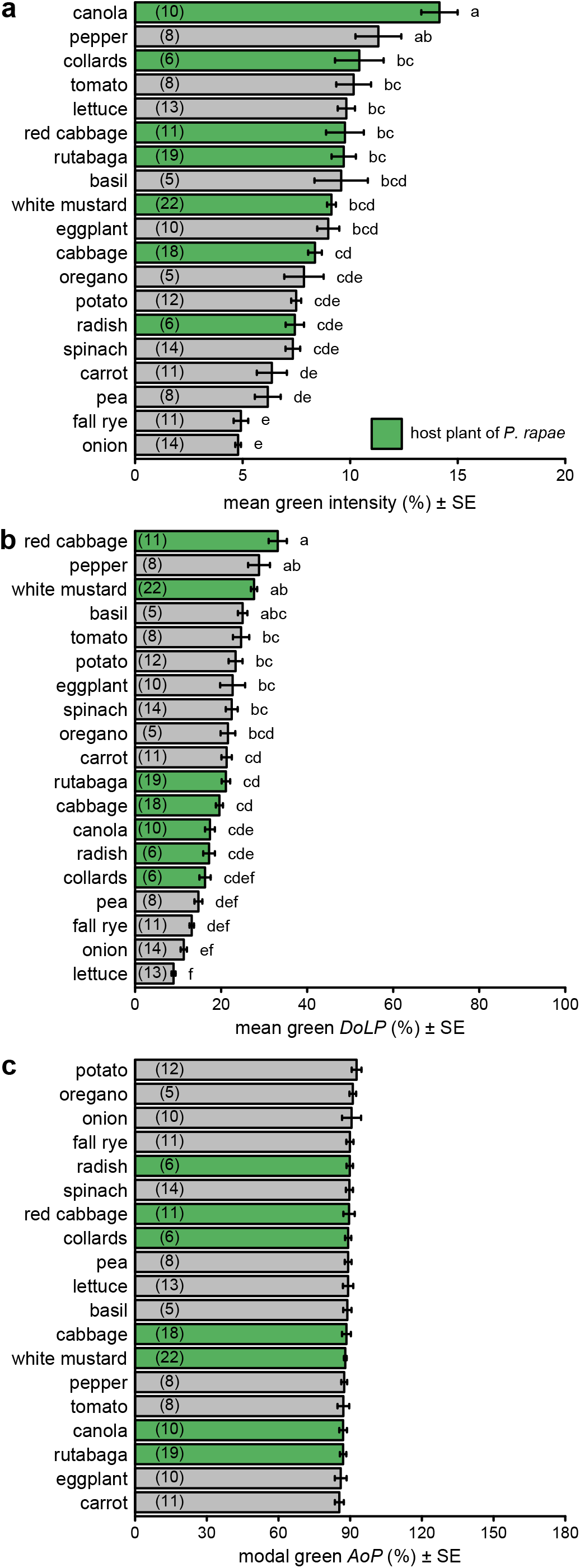
Comparison of intensity (**a**), degree of linear polarization (*DoLP*) (**b**), and axis of polarization (*AoP*) (**c**) among host plants (green bars) and non-host plants (grey bars) of *Pieris rapae*. These measurements used the green color band. Bars show mean or modal values with number of plants measured noted in parentheses in each bar. Bars with different letters differ statistically (*p*<0.05), as determined by a post-hoc Tukey test.

**Figure S5.**
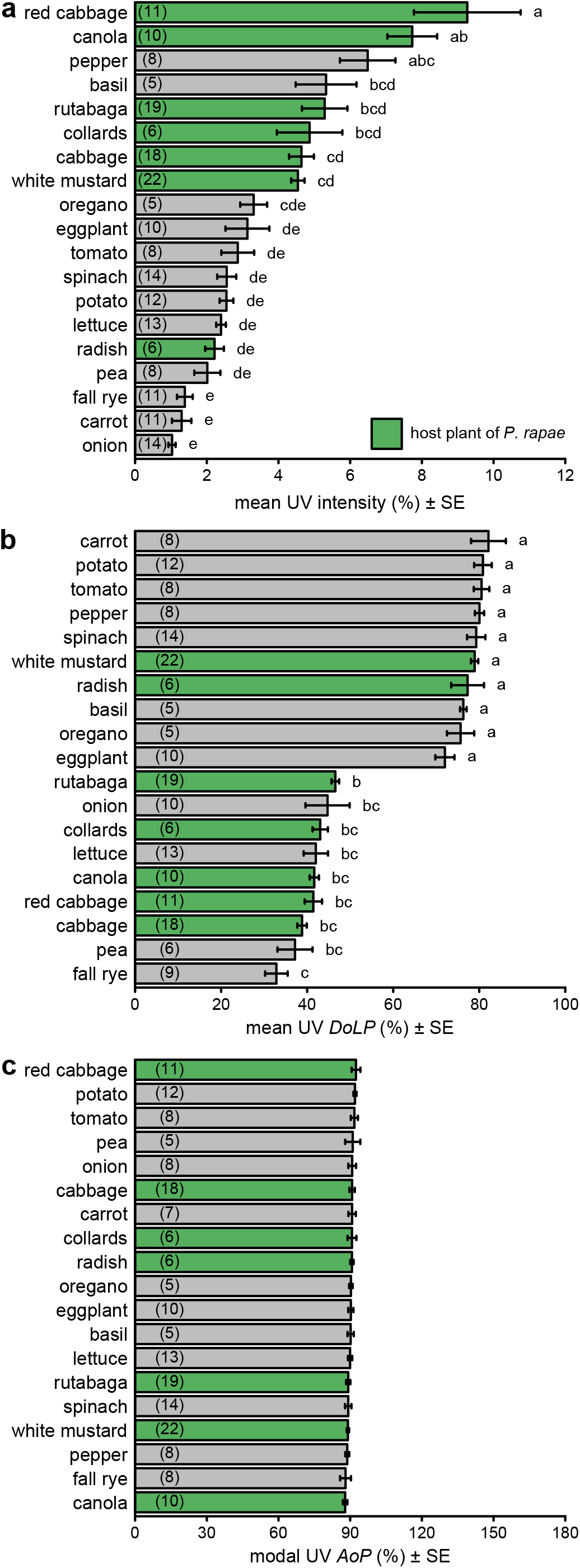
Comparison of intensity (**a**), degree of linear polarization (*DoLP*) (**b**), and axis of polarization (*AoP*) (**c**) among host plants (green bars) and non-host plants (grey bars) of *Pieris rapae*. These measurements used the UV color band. Bars show mean or modal values with number of plants measured noted in parentheses in each bar. Bars with different letters differ statistically (*p*<0.05), as determined by a post-hoc Tukey test.

**Figure S6.**
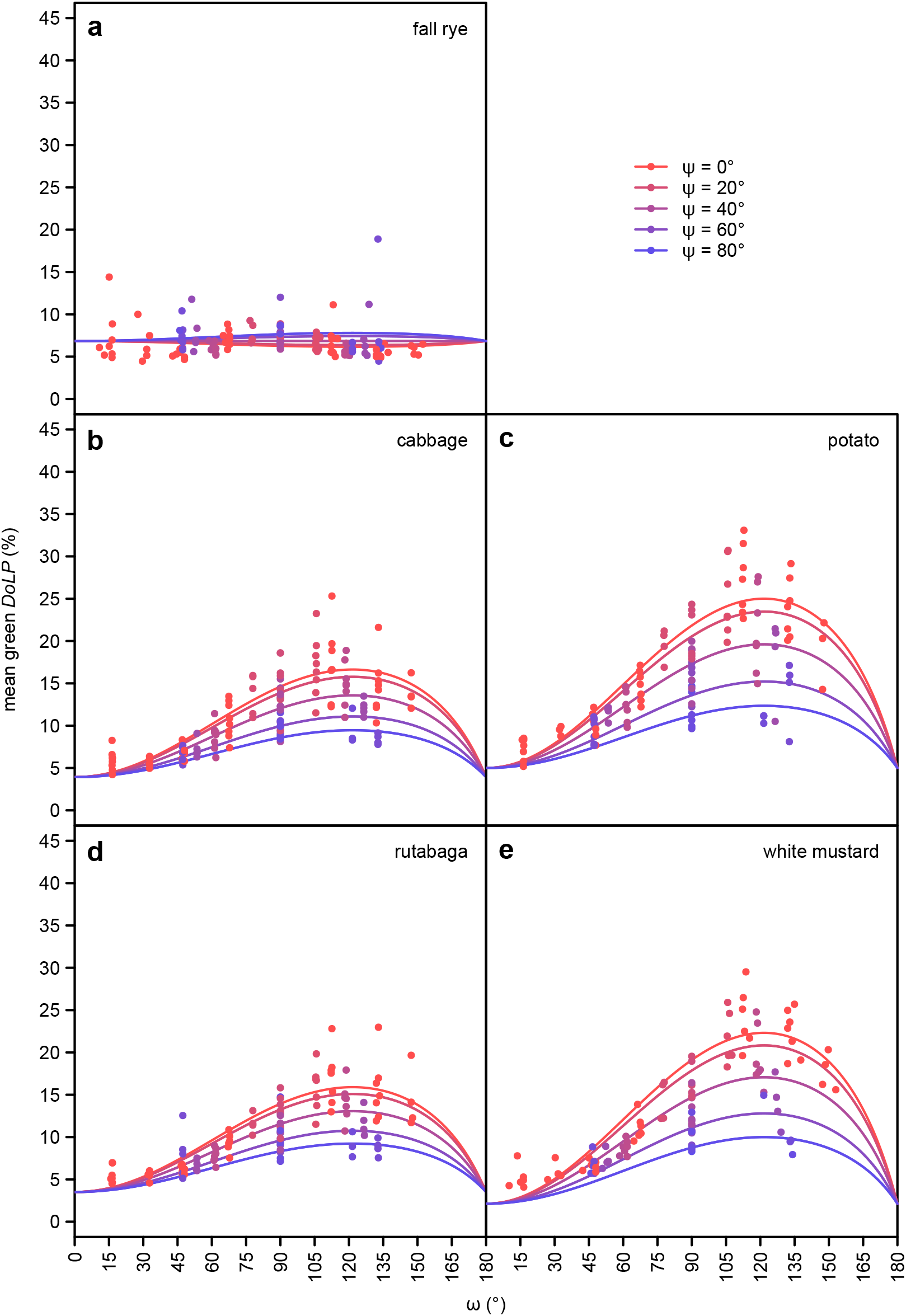
The effect of ω (angle between observer and light source with the plant at its vertex; see Fig. 1) and ψ (2-dimensional component of ω perpendicular to the plane passing through both the observer and the plant; see Fig. 1) on the mean degree of linear polarization (*DoLP*) of the green color band, as measured in five select plant species using photo polarimetry. Cabbage, rutabaga and white mustard are host plants of *Pieris rapae*.

**Figure S7.**
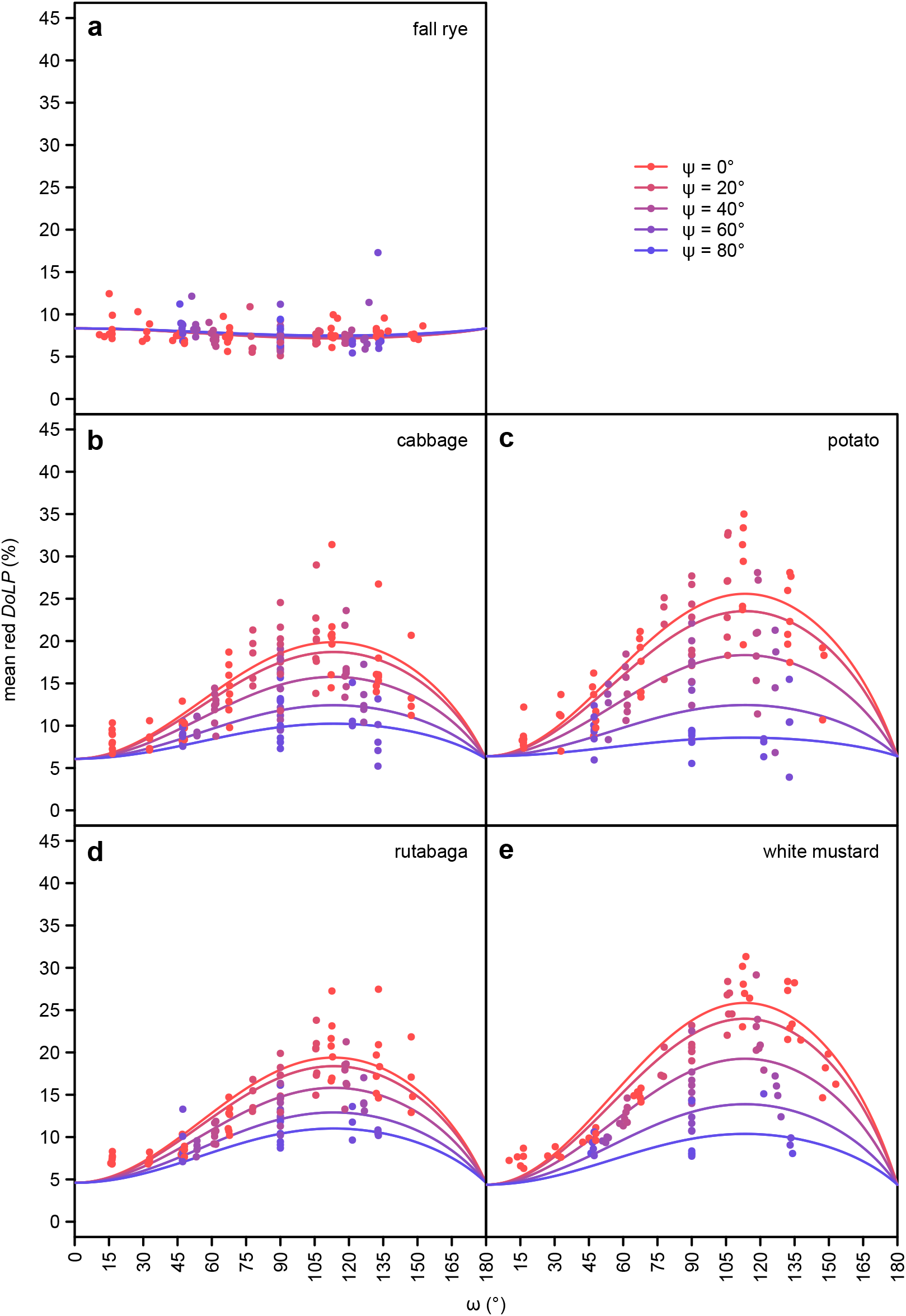
The effect of ω (angle between observer and light source with the plant at its vertex; see Fig. 1) and ψ (2-dimensional component of ω perpendicular to the plane passing through both the observer and the plant; see Fig. 1) on the mean degree of linear polarization (*DoLP*) of the blue color band, as measured in five select plant species using photo polarimetry. Cabbage, rutabaga and white mustard are host plants of *Pieris rapae*.

**Figure S8.**
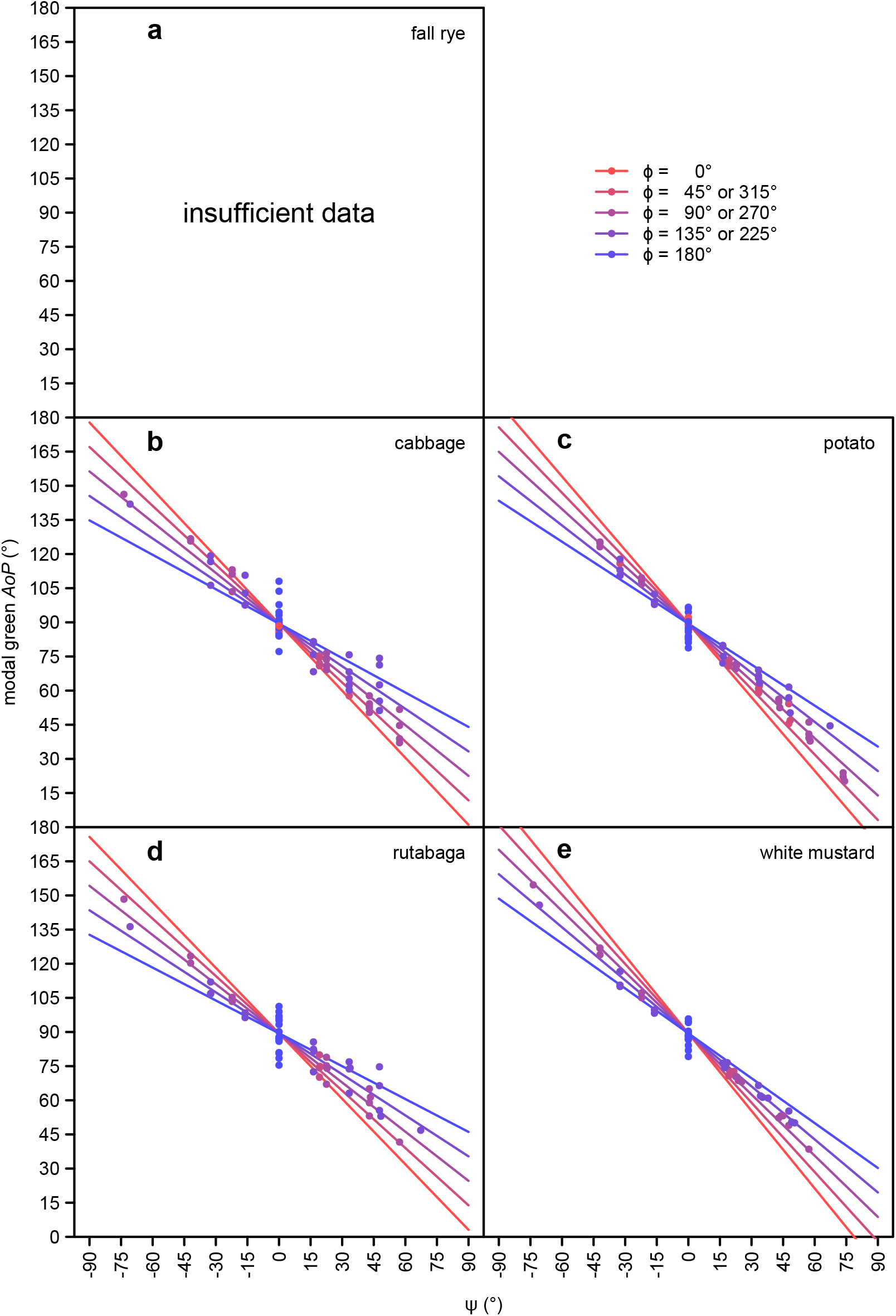
The effect of ψ (2-dimensional component of ω perpendicular to the plane passing through both the observer and plant; see Fig. 1) and ϕ (angle between the azimuth of the observer and the light source; see Fig. 1) on the modal axis of polarization (*AoP*) of the green color band, as measured in five select plant species using photo polarimetry. Cabbage, rutabaga and white mustard are host plants of *Pieris rapae*. Fall rye data were excluded from analyses due to an insufficient number of measurements meeting the inclusion criterion (>10% of pixels with a degree of linear polarization above 15%).

**Figure S9.**
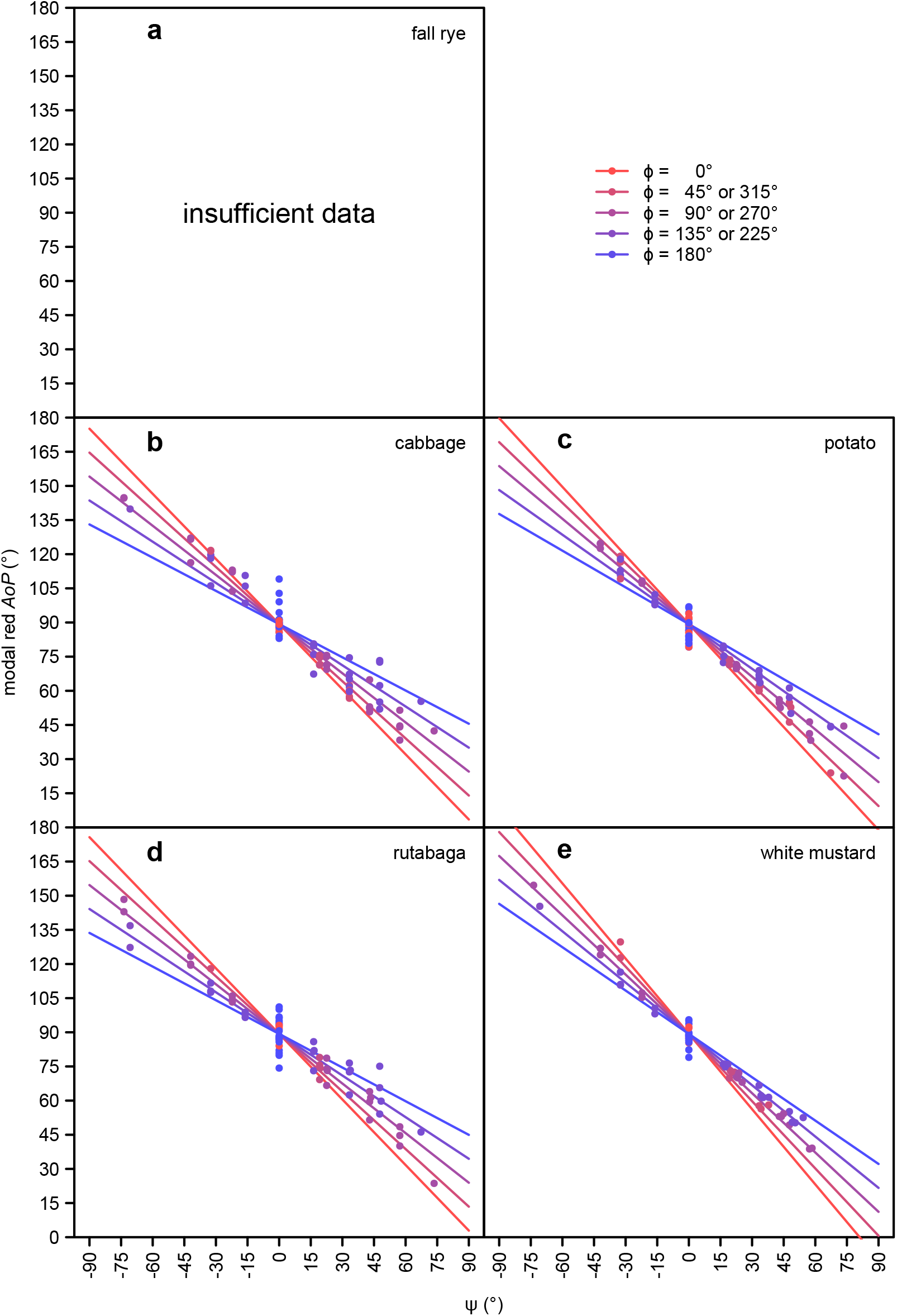
The effect of ψ (2-dimensional component of ω perpendicular to the plane passing through both the observer and plant; see Fig. 1) and ϕ (angle between the azimuth of the observer and the light source; see Fig. 1) on the modal axis of polarization (*AoP*) of the red color band, as measured in four select plant species using photo polarimetry. Cabbage, rutabaga and white mustard are host plants of *Pieris rapae*. Fall rye data were excluded from analyses due to an insufficient number of measurements meeting the inclusion criterion (>10% of pixels with a degree of linear polarization above 15%).

**Figure S10.**
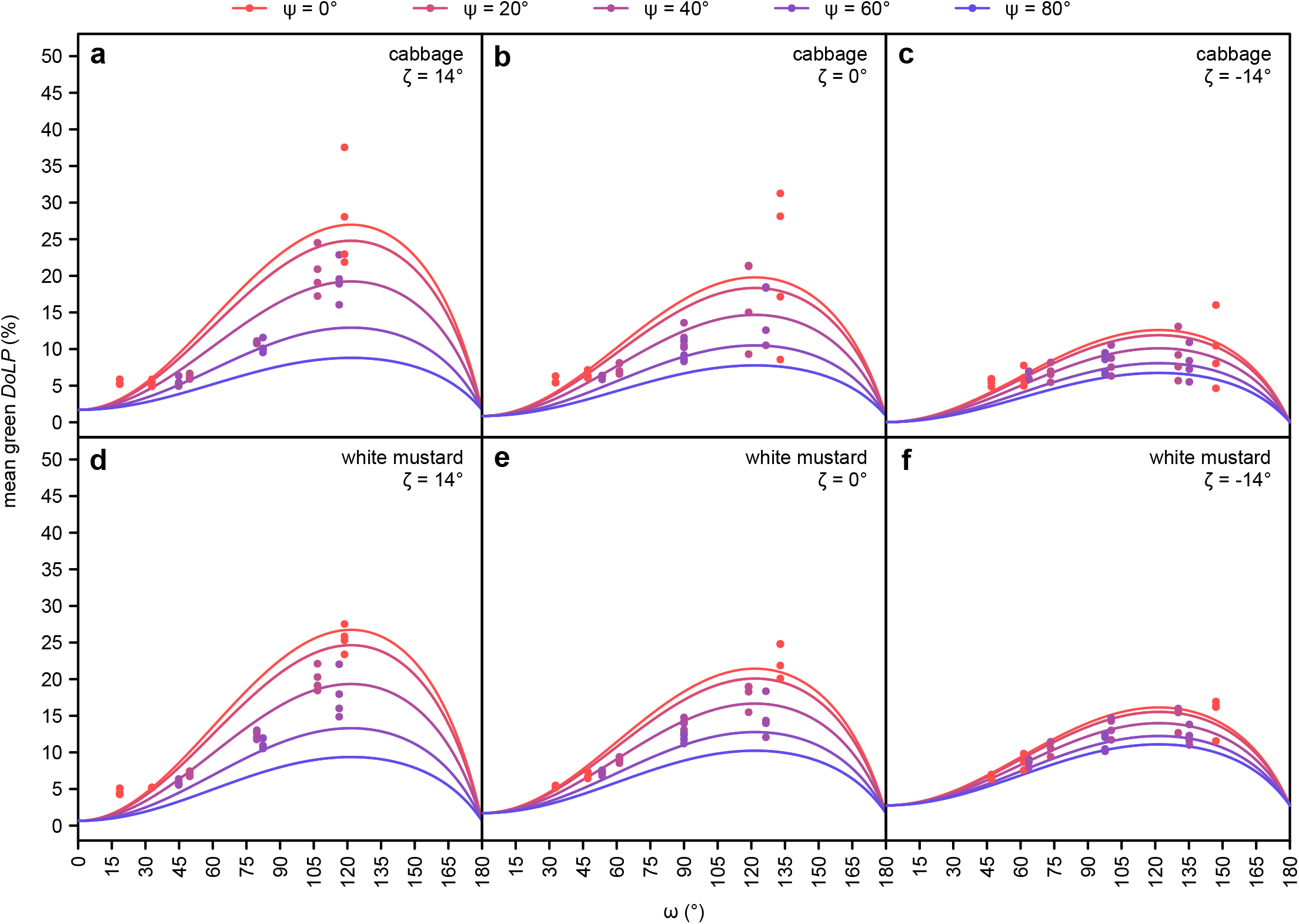
Additional effect of ζ (elevation of the observer; see Fig. 1) on the mean degree of linear polarization (*DoLP*) of the green color band, as measured in cabbage and white mustard (host plants of *Pieris rapae*) using photo polarimetry.

**Figure S11.**
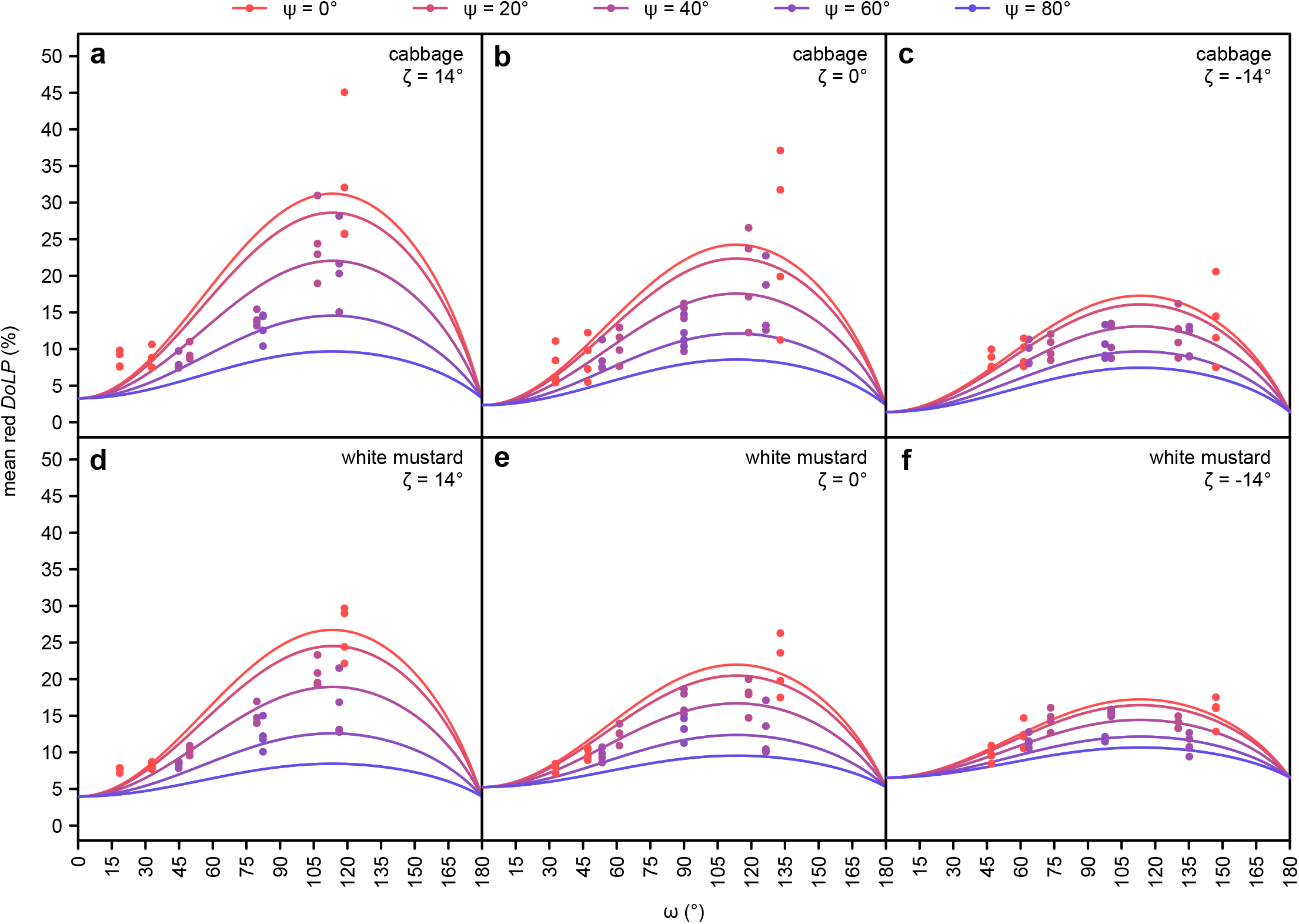
Additional effect of ζ (elevation of the observer; see Fig. 1) on the mean degree of linear polarization (*DoLP*) of the red color band, as measured in cabbage and white mustard (host plants of *Pieris rapae*) using photo polarimetry.

**Figure S12.**
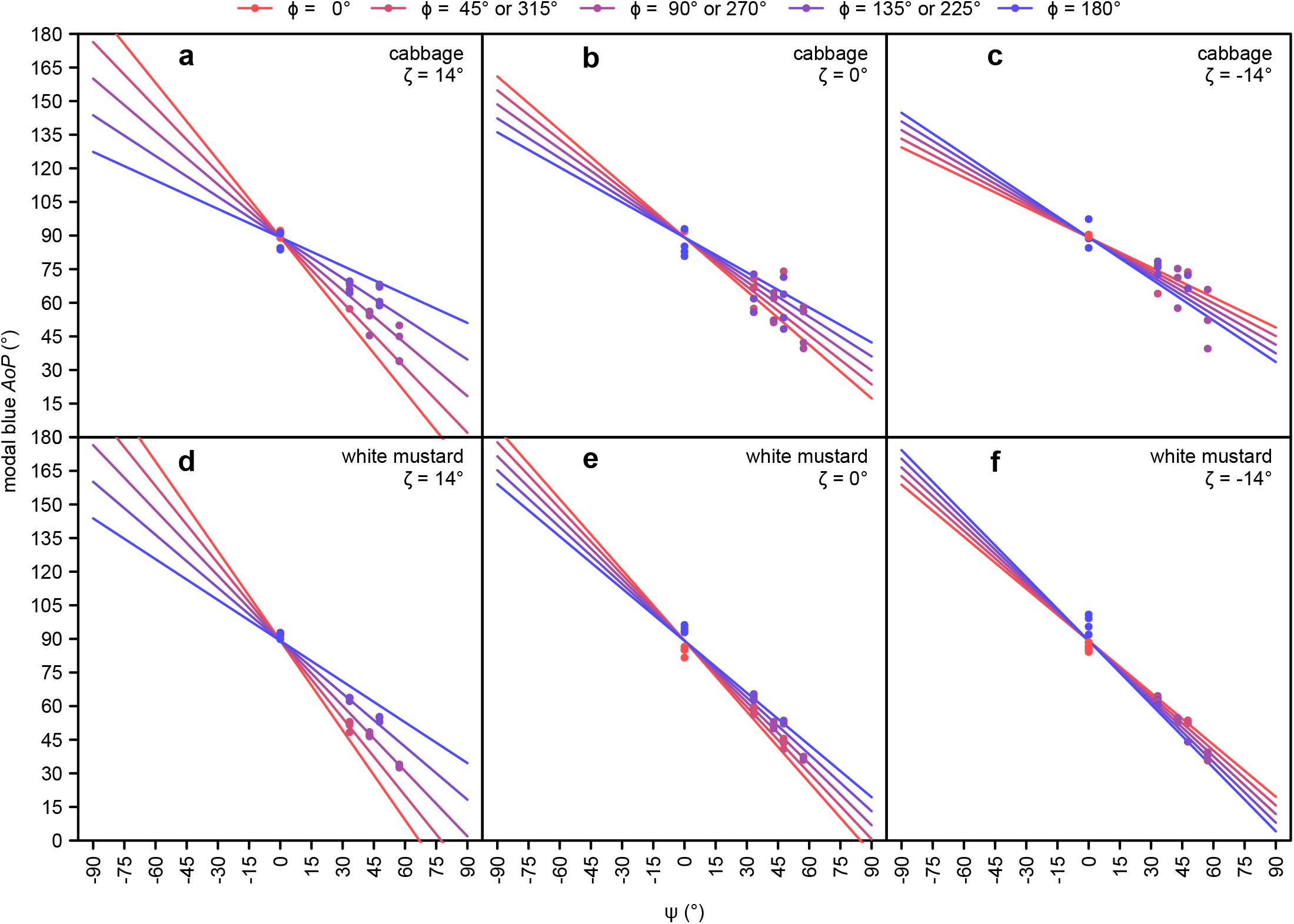
Additional effect of ζ (elevation of the observer; see Fig. 1) on the modal axis of polarization (*AoP*) of the blue color band, as measured in cabbage and white mustard (hosts of *Pieris rapae*) using photo polarimetry. Red and green color band data were excluded from analyses due to an insufficient number of measurements meeting the inclusion criterion (<10% of pixels with a degree of linear polarization above 15%).

**Figure S13.**
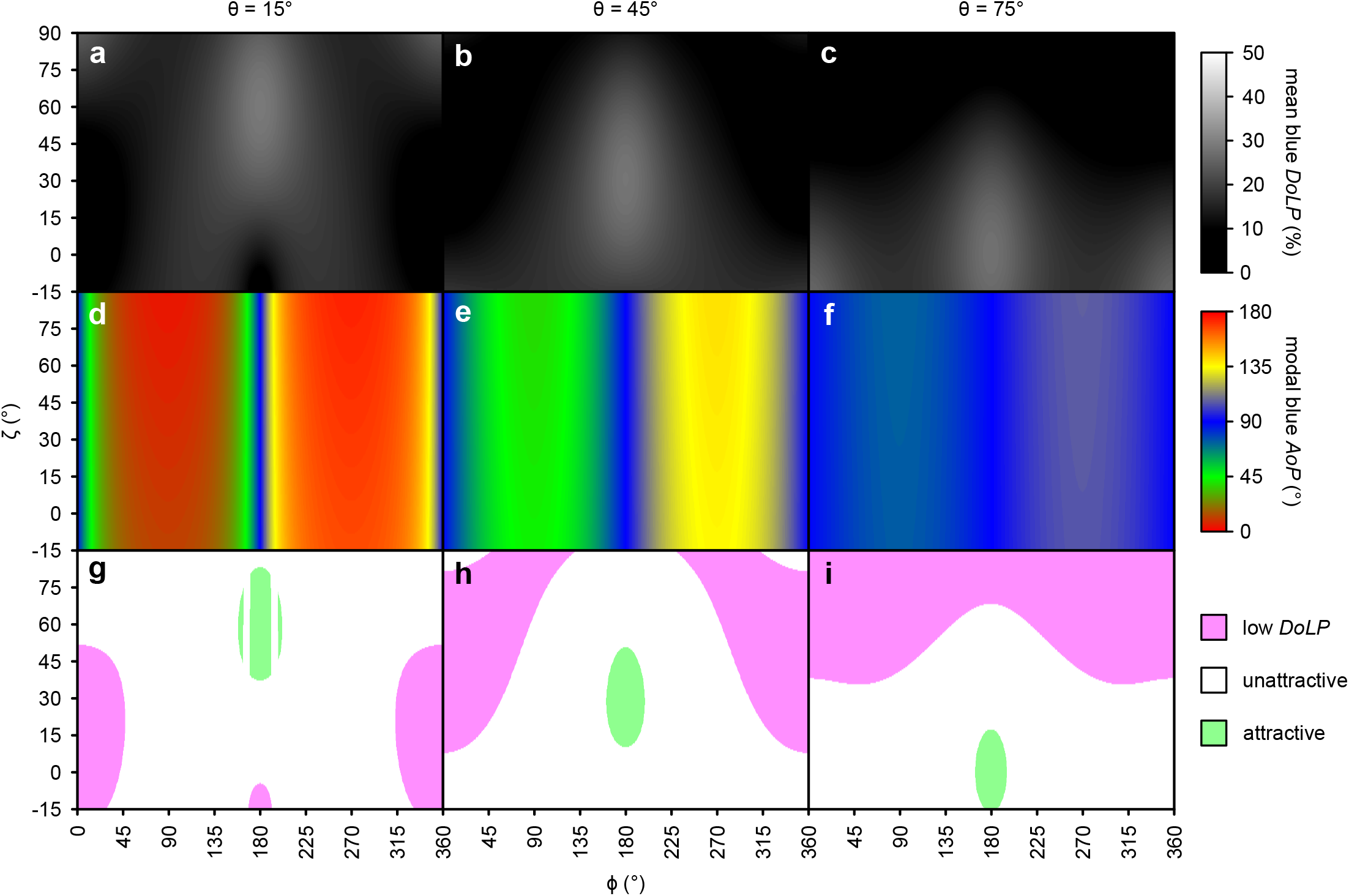
Effects of approach direction (angle between the azimuth of the observer and the light source (ɸ, see Fig. 1) and elevation of the observer (ζ; see Fig. 1) on the mean degree of linear polarization (*DoLP*) (**a-c**) and the modal axis of polarization (*AoP*) (**d-f**) of the blue color band of white mustard plants (host of *Pieris rapae*). Attractiveness of resulting polarization characteristics to *Pieris rapae* (**g-i**), based on a previous behavioral study (Blake et al. 2019). Approach trajectories resulting in attractive characteristics (*DoLP* = 26-36% and *AoP* = 0-38, 53-128 or 143-180°) and unattractive characteristics (*DoLP* = 10-26% or *AoP* = 38-53°, 128-143°) are shown in green and white, respectively, with pink indicating trajectories resulting in a moderately-attractive low *DoLP* (<10%). Higher *DoLP* (36-60%) would also be unattractive but were not predicted by these models. These effects changed with light source elevation (θ; see Fig. 1) which is shown at 15° (**a, d, g**), 45° (**b, e, h**) and 75° (**c, f, i**).

